# Fluorescent *in vivo* editing reporter (FIVER): A novel multispectral reporter of *in vivo* genome editing

**DOI:** 10.1101/2020.07.14.200170

**Authors:** Peter A. Tennant, Robert G. Foster, Daniel O. Dodd, Ieng Fong Sou, Fraser McPhie, Nicholas Younger, Laura C. Murphy, Matthew Pearson, Bertrand Vernay, Margaret A. Keighren, Peter Budd, Stephen L. Hart, Roly Megaw, Luke Boulter, Pleasantine Mill

## Abstract

Advances in genome editing technologies have created opportunities to treat rare genetic diseases, which are often overlooked in terms of therapeutic development. Nonetheless, substantial challenges remain: namely, achieving therapeutically beneficial levels and kinds of editing in the right cell type(s). Here we describe the development of FIVER (fluorescent *in vivo* editing reporter) — a modular toolkit for *in vivo* detection of genome editing with distinct fluorescent read-outs for non-homologous end-joining (NHEJ), homology-directed repair (HDR) and homology-independent targeted integration (HITI). We demonstrate that fluorescent outcomes reliably report genetic changes following editing with diverse genome editors in primary cells, organoids and *in vivo*. We show the potential of FIVER for high-throughput unbiased screens, from small molecule modulators of genome editing outcomes in primary cells through to genome-wide *in vivo* CRISPR cancer screens. Importantly, we demonstrate its *in vivo* application in postnatal organ systems of interest for genetic therapies — retina and liver. FIVER will broadly help expedite the development of therapeutic genome surgery for many genetic disorders.

## Introduction

The development of ever more precise and efficient genome editing technologies is revolutionising the ability to specifically and precisely alter the genome. Several clinical trials are currently underway using zinc finger nucleases (ZFNs), transcription activator-like effector nucleases (TALENs) and CRISPR/ Cas9 (clustered regularly interspaced short palindromic repeats (CRISPR)/CRISPR associated protein 9) based approaches for therapeutic targeted genome editing (1,2). The majority of these trials make use of *ex vivo* editing, however most genetic diseases would require somatic *in vivo* genome editing.

A major hurdle is the ability to efficiently monitor genome editing *in vivo*. Limited methods exist to track where, when and what types of editing outcomes occur *in vivo*, with most relying on next generation sequencing (NGS) to monitor changes at the DNA level (3–8). However, NGS technologies lack the spatial and temporal resolution needed to define which, and what proportion of cell types are edited in complex tissues. There is a need for simple, robust and cost-effective systems allowing for rapid detection of genome editing *in vivo*. Genetically-encoded fluorescent reporters offer one potential solution, allowing a rapid visual read-out at both a cellular and organismal level which can be easily quantified both by microscopy and flow cytometry.

All genome editing methods rely on the cell’s own machinery to repair the targeted DNA double strand breaks (DSBs). Broadly speaking, they use one of two major pathways (9): non-homologous end-joining (NHEJ), often leading to small insertions or deletions (indels); and when a template is available, homology directed repair (HDR), resulting in precise correction of disease-causing mutations. Several fluorescence-based reporter systems for monitoring the outcomes of genome editing have been described (10–17). However, these are predominantly *in vitro* reporters, relying on transiently transfected constructs or stable cell lines, or where available *in vivo* are limited to the detection of NHEJ events (14,15). *In vitro*, these reporters have been useful to expedite discovery of small molecule modifiers of genome editing outcomes. However, efficiently expanding their use *in vivo* towards precisely controlled genome editing, or ‘genome surgery’, in target cells requires a different approach.

To address these issues, we have developed a novel fluorescent *in vivo* editing reporter (FIVER) mouse model, which generates a visible, quantifiable fluorescence read-out of different editing outcomes in real time with single cell resolution. This allows direct visualisation of NHEJ, HDR and homology-independent targeted integration (HITI) based (18) editing by distinct fluorescent outcomes. FIVER allows rapid side-by-side evaluation of different delivery methods (i.e., viral or non-viral) and payloads by altering choice of genome editors or repair sequences used. It also lends itself to screening small molecule modifiers of DNA repair pathways which might promote desired editing outcomes *in vivo*.

Importantly, we have developed the FIVER genome editing toolkit to be used in the widely available *mTmG* Cre-reporter mouse model (19) to facilitate rapid uptake by the community. Here, we describe an *in vivo* fluorescent genome editing reporter, which is the first that is able to monitor a range of genome editing outcomes, both templated (HDR or HITI) and non-templated (NHEJ), via multispectral readouts of these events throughout the entire lifespan of the animal and their fates in complex tissues.

## Results

### Development of a tricolour luorescent reporter for CRISPR-based genome editing

In order to design a responsive and reproducible *in vivo* genome editing reporter, we set out to develop a modular system that could be used for *in vivo*, *ex vivo*, and primary cell line genome editing in mice. To facilitate widespread uptake by the scientific community, we repurposed the previously described *mTmG* Cre-mediated recombination reporter mouse (19), in which a ubiquitous CAG promoter drives expression of a floxed membrane-tagged tdTomato gene followed by a strong transcriptional stop element at the *Rosa26* locus, which is in turn followed by a membrane-tagged EGFP. Targeting genome editing tools to create DSBs near both loxP sites flanking the tdTomato gene should yield results analogous to Cre-mediated recombination, such that a shift in fluorescence, from tdTomato to EGFP, would reflect genome editing activity. Henceforth, we will refer to heterozygous *mTmG* animals as FIVER for clarity.

We identified *Streptococcus pyogenes* Cas9 (SpCas9) guide-RNA (gRNA) target sites in a conserved region adjacent to both loxP sites flanking the tdTomato coding sequence. We selected the top scoring (in terms of predicted off target profile) SpCas9 gRNAs targeting both the antisense and sense strands — T1 and T2, respectively (**Figure 1A**). In primary mouse embryonic fibroblasts (MEFs), both guides result in excision of the intervening tdTomato coding sequence; we have focused on T1. Repair of this lesion results in distinct fluorescent changes depending on the repair pathway employed. When no repair template is supplied, the lesion is repaired via NHEJ which can allow expression of the downstream membrane-tagged EGFP (mEGFP). In addition, when asynchronous cleavage occurs, indels at the upstream site can lead to loss of tdTomato expression, but not removal of the tdTomato cassette resulting in total loss of fluorescence (**Figure 1A**). By simultaneously supplying an exogenous repair template, the membrane tag of EGFP can be exchanged for a nuclear localised signal from histone H2B, resulting in expression of a nuclear EGFP (nEGFP) (**Figure 1 and figure supplement 1**). H2B provided a more robust nuclear signal than canonical NLS sequences (**Figure 1-figure supplement 1B**) and thus was ideal for automated detection, across cell types and cell cycle stages, and was used for all subsequent experiments.

**Figure 1.**
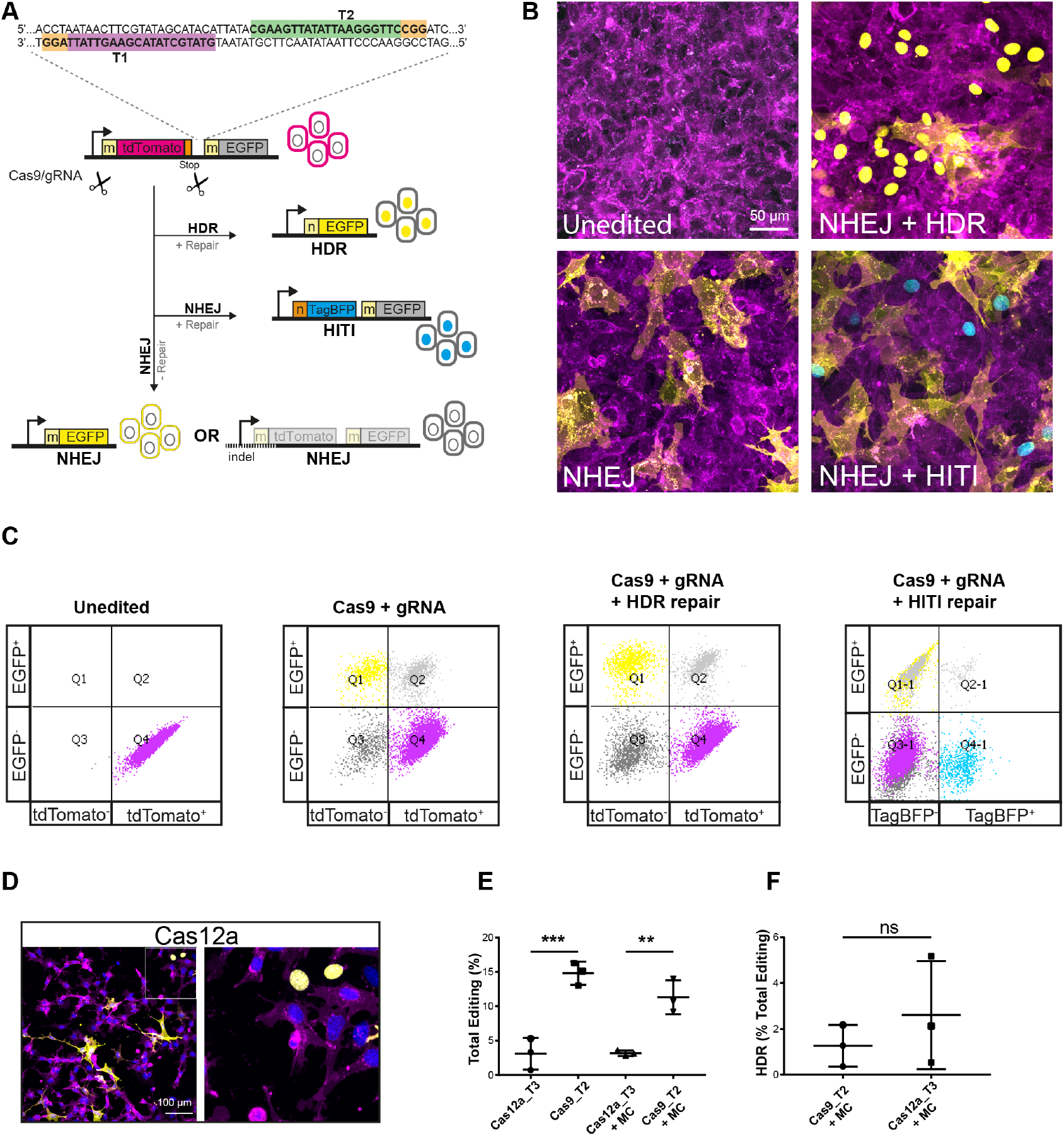
Overview of luorescent *in vivo* editing reporter (FIVER) system. (A) Schematic of FIVER system. We identified conserved gRNA sites on both the sense (T2; green box) and antisense (T1; purple box) strands flanking the tdTomato cassette within the FIVER locus (PAM sites indicated by orange boxes). Here membrane-tagged tdTomato is expressed by every cell. When CRISPR machinery and either T1 or T2 gRNA are provided, the tdTomato cassette is excised. Without the provision of an exogenous repair template non-homologous end joining (NHEJ) repair is employed to repair the lesion, allowing expression of downstream membrane-tagged EGFP, observed with a shift from membrane tdTomato (mtdTomato) to membrane EGFP fluorescence (mEGFP). Alternatively, asynchronous cleavage and/or larger indels (dotted line) can cause disruption of the tdTomato resulting in loss of all fluorescence. If a template containing homology to the locus is provided, the lesion can be repaired by homology directed repair (HDR), in our system this replaces the membrane tag of the downstream EGFP for a nuclear tag (H2B) resulting in a shift from mtdTomato to nuclear EGFP (nEGFP) fluorescence. Finally, if a homology-independent targeted integration (HITI) repair template is provided, then NHEJ can be employed to knock in a membrane-tagged TagBFP construct, resulting in a shift from mtdTomato to nuclear TagBFP (nTagBFP) fluorescence. m = MARCKS membrane tag, n = H2B nuclear localisation signal. (B) Representative confocal images of mouse embryonic fibroblast (MEF) lines derived from FIVER mice and edited with and without repair constructs. Images are maximum intensity projections from z-stacks. (C) Representative flow cytometry plots following editing in MEF lines. All editing outcomes can be observed, however nEGFP and mEGFP are indistinguishable by this method (see **Figure 1-figure supplement 2**). FACS was carried out 5 days post transfection. (D) Representative confocal images of MEFs edited using AsCas12a machinery with T3 gRNA. Nuclei are stained with Hoechst. (E) Editing in MEF lines using Cas9 is significantly more efficient than using AsCas12a (*p* <0.001; one-way ANOVA with Tukey’s multiple comparison), n = 10,000 single cells, N = 3 technical replicates. (F) There is no significant difference in the ability of SpCas9 and AsCas12a to drive HDR in MEF lines using minicircle (MC) delivery of repair constructs (*p* = 0.257; unpaired t-test), n > 6,000 cells, N = 3 technical replicates. **Figure 1–Figure supplement 1. Overview of luorescent *in vivo* editing reporter (FIVER) system**. **Figure 1–Figure supplement 2. Overview of luorescent *in vivo* editing reporter (FIVER) system**.

In addition to reporting on NHEJ or HDR, we included a read-out for HITI, an NHEJ-based method for specifically altering the genome (18). Cas9-induced DSBs in both the target locus and in the delivered repair plasmid allow the fragment generated during editing to integrate into the genomic locus without the need for sequence homology. As this method relies on the NHEJ pathway, it can occur at any point during the cell cycle and in terminally differentiated cells (20,21). HITI has great therapeutic potential in that an exogenous cDNA could be introduced under endogenous transcriptional control. We designed a HITI donor construct consisting of a nuclear-localised TagBFP followed by a strong stop sequence. This read-out is both spectrally distinct from tdTomato and EGFP and spatially distinct from the membrane localisation of the reporter. Following excision of tdTomato, HITI repair leads to TagBFP knock-in. This can be visualised as a switch from membrane tdTomato fluorescence (mtdTomato) to nuclear localised TagBFP fluorescence (nTagBFP) (**Figure 1A-C**).

To test the system, we generated immortalised MEF lines from FIVER mice and transiently transfected them with ribonucleoprotein (RNP) comprised of SpCas9 protein complexed with either T1 or T2 gRNAs. Confocal imaging and flow cytometry confirmed transition from mtdTomato to mEGFP, accounting for approximately 30% of events, indicative of NHEJ repair following excision of the tdTomato cassette (**Figure 1B-C**). In addition, there was a total loss of fluorescence following CRISPR/Cas activity in approximately 30% of cells, due to larger deletions or imperfect repair which truncated the fluorophore or altered the reading frame. In some instances (particularly in immortalised MEF lines, accounting for approximately 10% of events, but not *in vivo*) we also observed a tdTomato^+^ /EGFP^+^ population following editing; primarily observed using flow cytometry. As a result, we took the total of tdTomato^−^/EGFP^+^, tdTomato^+^/EGFP^+^ and tdTomato^−^/EGFP^−^ populations to represent overall levels of editing.

To assess HDR pathways, we constructed both single- and double-stranded repair templates, containing homology arms of various lengths (35 bp to 780 bp) and have focused on ~700 bp arms flanking an H2B nuclear localisation signal encoded on a minicircle vector (**Figure 1–figure supplement 1A**) which initially gave the highest and most consistent rates of repair (**Figure 1–figure supplement 1C**). Following co-delivery of this construct (MC.HDR) with CRISPR/Cas machinery, nEGFP fluorescence could be observed (**Figure 1B**). The edited cells were also subjected to flow cytometric analysis (**Figure 1C**). A shift in fluorescent profile was observed following editing, however, nEGFP and mEGFP expression were not distinguishable by intensity using standard flow cytometry (**Figure 1–figure supplement 2**), necessitating an image analysis-based approach to quantify HDR, as described later.

The method of delivering editing machinery can impact editing outcomes and will vary depending on application (22). To address this, we have built a toolkit to allow delivery of CRISPR components and repair constructs in various forms (RNP, plasmid or minicircle) either by non-viral methods (i.e., nucleofection, lipid nanoparticles and hydrodynamic injection) or virally (i.e., lentivirus and adeno-associated virus).

As the FIVER system reports on DSB-repair outcomes, we postulated that any site specific nuclease generating DSBs could be employed. While the bulk of work has focused on SpCas9, differences in nuclease size, types of ends generated and availability of specific PAM motifs close to the target may warrant use of a range of genome editors. Therefore, we designed gRNAs for use with *Staphylococcus aureus* Cas9 (SaCas9) and *Acidaminococcus sp.* Cas12a (AsCas12a) (previously Cpf1) to target the same conserved region flanking tdTomato. We assayed the activity of AsCas12a and demonstrated the ability of FIVER to report its editing outcomes (**Figure 1D-F**). In addition, potential target sites for TALENs are listed in **Supplementary Table 1**. In summary, FIVER is a robust fluorescent reporter of genome editing events for a range of DSB-inducing genome editors.

### DNA sequencing confirms fidelity of luorescent read-outs relecting underlying genetic changes

In order to confirm the reliability of the fluorescent read-out and identify the origin of the double positive tdTomato^+^/EGFP^+^ population, we carried out deep sequencing on edited cell populations. MEFs transfected with SpCas9-RNP-based editing reagents and minicircle HDR template (MC.HDR) were sorted into four populations — tdTomato^+^/EGFP^−^ (unedited), tdTomato−/EGFP^+^ (NHEJ and HDR), tdTomato^−^/EGFP^−^ (NHEJ) and tdTomato^+^/EGFP^+^ (unexpected outcome) (**Figure 2–figure supplement 1C**). The reporter locus was amplified from genomic DNA isolated from each population by PCR (**Figure 2A**) and the amplicons sequenced on both the Ion Torrent and MinION sequencing platforms. This combinatorial approach allowed us take advantage of the longer MinION reads for detection of larger structural changes, while retaining the greater base calling accuracy of Ion Torrent reads.

**Figure 2.**
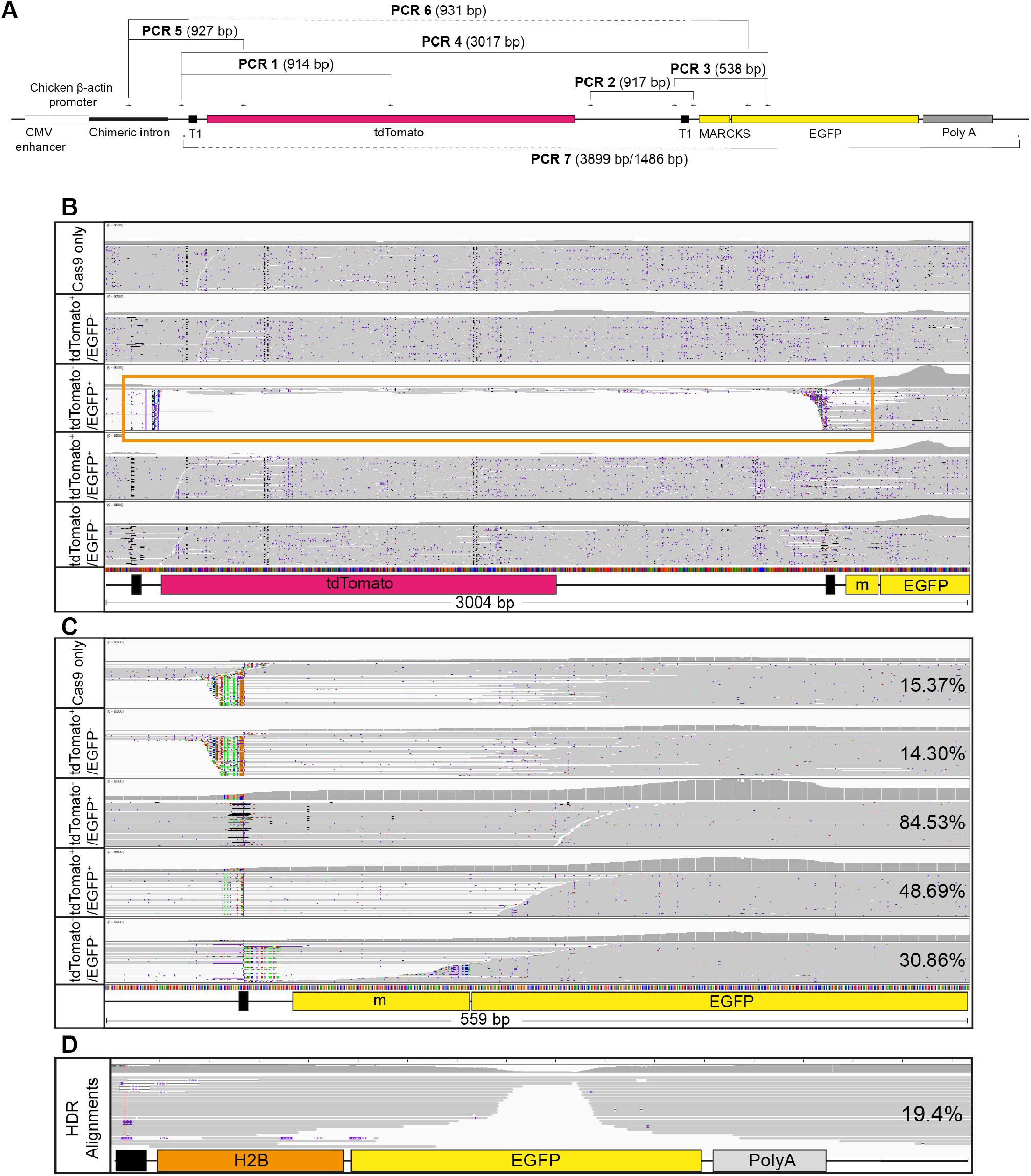
Deep sequencing confirms editing outcomes observed by FIVER. (A) Overview of FIVER locus, with primers and PCRs used for sequencing indicated. (B) Ion Torrent reads from PCR product 4 mapped to the locus for each sorted population of cells. Orange box indicates loss of tdTomato cassette in tdTomato-/EGFP^+^ population. Filled black boxes indicate T1 target sites, m = MARCKS membrane tag. (C) Ion Torrent reads from PCR product 4 mapped to the predicted NHEJ product (i.e., removal of tdTomato) for each sorted population of cells. Filled black box indicates T1 target region, m = MARCKS membrane tag. Percentage of reads correctly aligned for each population are indicated. (D) Reads from TOPO cloned and sequenced samples from the tdTomato^−^/EGFP^+^ population (PCR 7), mapped against the predicted HDR outcome. m = MARCKS membrane tag. The percentage of reads which align are indicated. **Figure 2–Figure supplement 1. Deep sequencing confirms editing outcomes observed by FIVER**.

We first employed variant calling to map the Ion Torrent reads to a predicted reference sequence based on anticipated outcomes (**Figure 2B-C**). Using this approach, >90% of reads within the tdTomato^+^/EGFP^−^ (unedited) population aligned with the reference sequence. However, 21.28% of reads across the upstream gRNA target site demonstrated indels and a further 7.67% of reads harboured indels at the downstream gRNA site, compared to 1.5% and 1.47%, respectively for the untreated (Cas9 only) population. This suggests low levels of cleavage and subsequent repair by NHEJ at the two sites that were under-reported by FIVER, with a maximum false negative rate of 26.48%. However, individual Ion Torrent reads are not long enough to cover both gRNA sites, meaning it is not possible to confirm if one or both sites contained indels in individual cells. For the tdTomato^−^/EGFP^+^ (NHEJ and HDR) population 84.53% of reads align to the expected NHEJ repair product, that is, complete removal of the tdTomato cassette between the two gRNA sites (**Figure 2C**). In contrast, only 15.37% and 14.3% of reads from Cas9 only and tdTomato^+^/EGFP^−^ (unedited) populations, respectively, aligned to the predicted NHEJ repair product (**Figure 2C**). However, these predominantly align across the EGFP gene and not the repair junction (**Figure 2C**). This suggests a high accuracy of the mEGFP readout.

Using *de novo* genome assembly, the MinION reads were successfully assembled in order to form the major sequences present within the input. When aligned to the reference, sequences from the Cas9 only control were assembled with very little error (**Figure 2–figure supplement 1A**). In addition, three sequences assembled from the tdTomato^−^/EGFP^+^ (NHEJ and HDR) population were all lacking tdTomato, confirming fidelity of this readout for NHEJ. Interestingly, the tdTomato^+^/EGFP^+^ double positive population consists of a mixture of sequences with (88%) and without (12%) tdTomato, confirming that this population results from editing at the locus. The tdTomato^−^/EGFP^−^ (NHEJ) population also appeared to be a mixture, with some sequences missing segments of tdTomato; which could explain their loss of fluorescence (**Figure 2–figure supplement 1A**).

To confirm the origins of the tdTomato^−^/EGFP^−^ (NHEJ) double negative population, we generated new PCR primers which anneal within the CAG promoter (**Figure 2A**, PCR 5 and 6) to capture larger deletions. This revealed that almost all tdTomato^−^/EGFP^−^ double-negative cells harbour large deletions that extend into the EGFP sequence and/or the promoter, anticipated to cause a total loss of fluorescence (**Figure 2–figure supplement 1B**). Taken together with the MinION data, this population results from larger indels, either with or without loss of tdTomato, confirming that this population is also the result of NHEJ repair.

To investigate HDR, targeted resequencing of the tdTomato^−^/EGFP^+^ (NHEJ and HDR) population was carried out. Primers spanning the entire locus were used to ensure that the long-lived minicircle donor template was not erroneously amplified (23,24) (**Figure 2A**, PCR 7). Of the tdTomato^−^/EGFP^+^ population, 19.4% of reads aligned to the predicted HDR sequence containing integrated H2B (**Figure 2D**). Given the rate of total editing here (**Figure 2–figure supplement 1C**), this means approximately 1.32% of total cells underwent HDR, consistent with the range of HDR efficiency we have previously observed in MEFs (**Figure 1–figure supplement 1B**) and a similar proportion to that described in the literature (25–27). This suggests that observed nEGFP is consistent with changes at the DNA level.

### Rapid transitions in luorescence upon genome editing

To determine the dynamics of the fluorescence transitions upon genome editing, we performed time-lapse imaging of primary MEFs following transfection with plasmid derived CRISPR, with and without minicircle repair constructs (MC.HDR or MC.HITI) (**Figure 3A and Figure 3–video 1**). For each condition, 30 random fields were imaged and edited cells identified based on final fluorescence. Mean intensities for each channel were calculated for each time point using the manual tracking Fiji plugin (**Figure 3B**).

**Figure 3.**
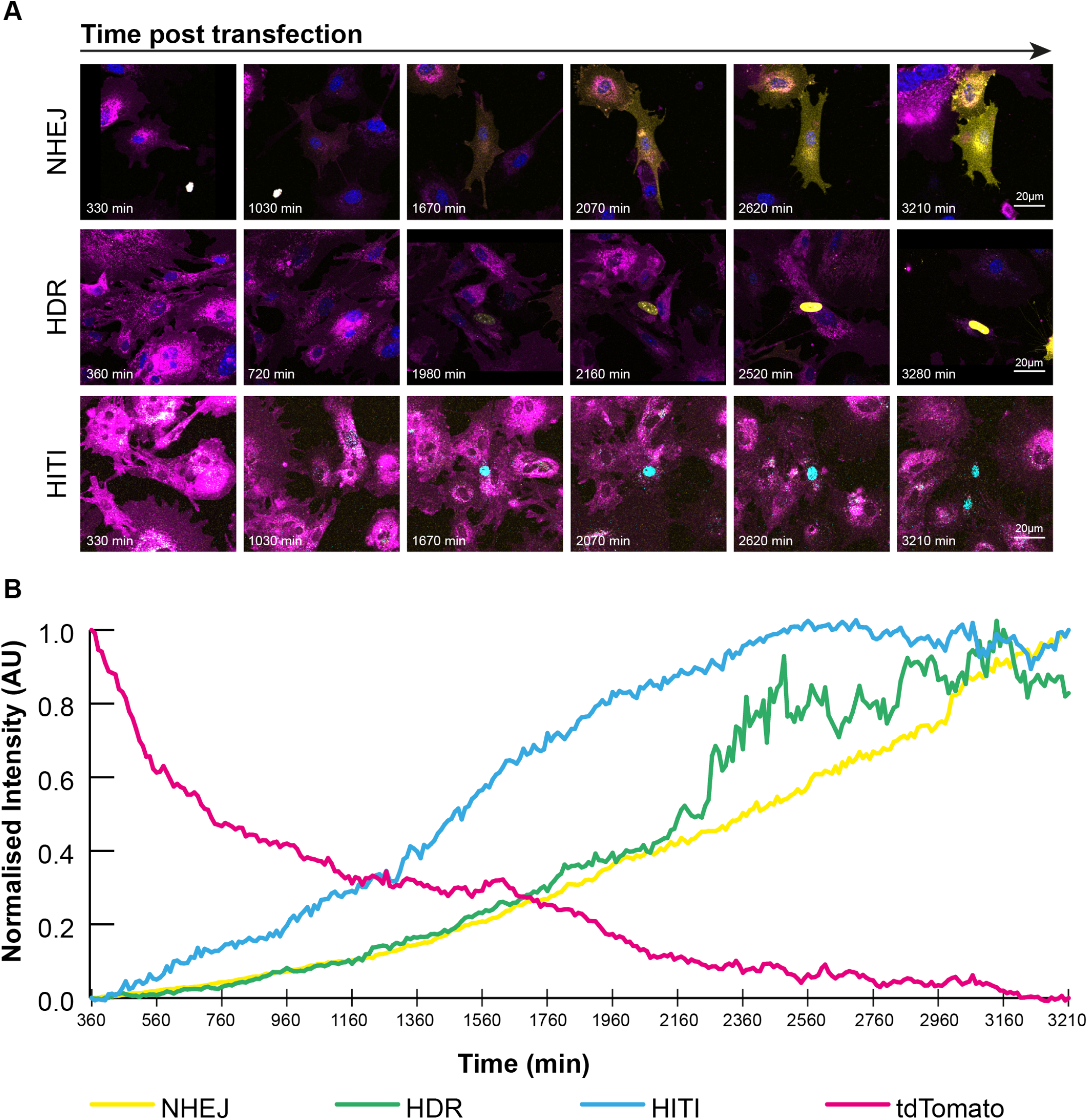
Rapid transition in luorescent signal following editing in FIVER MEFs. FIVER MEFs were nucleofected with plasmid and minicircle based CRISPR components (pX330-T1, MC.HDR and MC.HITI), then imaged at 30 random points per well in 6-well dishes every 10 min for 48 hours. Edited cells were analysed using the manual tracking plugin for ImageJ. (A) Representative cropped confocal images from time lapses of tracked cells, single z-slices. For NHEJ and HDR samples, nuclei are stained with Hoechst. (B) Means of normalised fluorescence intensity of tracked cells over time, n = 6 HDR, n = 26 NHEJ, n = 21 HITI and n = 53 tdTomato. For full time course see **Figure 3–video 1**. **Figure 3–video 1. Rapid transition in luorescent signal following editing in FIVER MEFs**. FIVER MEFs were nucleofected with plasmid and minicircle based CRISPR components (pX330-T1, MC.HDR and MC.HITI), then imaged at 30 random points per well in 6-well dishes every 10 min for 48 hours. Edited cells were analysed using the manual tracking plugin for ImageJ. Videos show full time lapse for each condition represented in **Figure 3A**. (A) Tracking of NHEJ edited cell. (B) Tracking of HDR edited cell. (C) Tracking of HITI edited cell. Scale bar 20 *μ*m.

In all edited cells, mtdTomato fluorescence rapidly decreased (magenta line, **Figure 3B**). In the case of NHEJ, this signal was concurrently replaced with mEGFP fluorescence, increasing gradually in mean intensity over time (yellow line, **Figure 3B**). In the case of HDR, nEGFP accumulates gradually before rapidly increasing in intensity, then plateauing (green line, **Figure 3B**). Similarly, for HITI editing, nTagBFP accumulates gradually at first, before rapidly increasing then plateauing approximately 40 hours post transfection (blue line, **Figure 3B**). In all cases, the switch in fluorescence occurs rapidly and is complete by 48 hours post-transfection (**Figure 3–video 1**).

### Screening small molecule modulators of genome editing outcome with FIVER

One of the limitations of genome editing as a therapeutic tool is its dependence on endogenous DNA repair pathways to resolve targeted nicks, cuts and/or breaks generated by the nucleases. The reliance on HDR to generate specific changes in the genomes of mammalian somatic cells, where this is not the dominant DNA repair pathway (28), has led to the search for methods to manipulate repair mechanism choice. This includes the identification of small molecules to bias outcomes towards precise repair by stimulating HDR as well as inhibiting NHEJ. However, it remains largely unknown whether all cell types will respond similarly in resolving genome edited DSBs and whether there are cell-type-specific effects of these small molecules.

Three main classes of small molecule have been shown to be effective at increasing the eff-ciency of HDR: (1) inhibitors of NHEJ (25,29–31); (2) enhancers of the HDR pathway (32–34); and (3) molecules of unknown mechanism(s) (27). FIVER cells are ideal for unbiased screening of compounds as image acquisition and analysis can be done in an automated (and blinded) manner and at scale. Initially, we tested five compounds which had been shown previously to increase the efficiency of CRISPR-based HDR, whose mechanisms of action are summarised in **Figure 4A**, to see if effects could be recapitulated in our FIVER MEF lines. Two of these disrupt NHEJ: NU7441, an inhibitor of DNA-dependent protein kinase catalytic subunit (DNA-PKcs) (31), and an inhibitor of DNA Ligase IV named Scr7 (29). RS-1, an activator of the homologous recombination protein Rad51 (33), has been reported to increase HDR efficiency in response to CRISPR-induced DNA damage. We also tested two molecules identified using a blind screening method for molecules which improved the efficiency of HDR in CRISPR edited cells (27), L755,501, an agonist of the *β*3 adrenergic receptor (35), and Brefeldin-A (Brf-A), an inhibitor of ADP ribosylation factor 1 (36).

**Figure 4.**
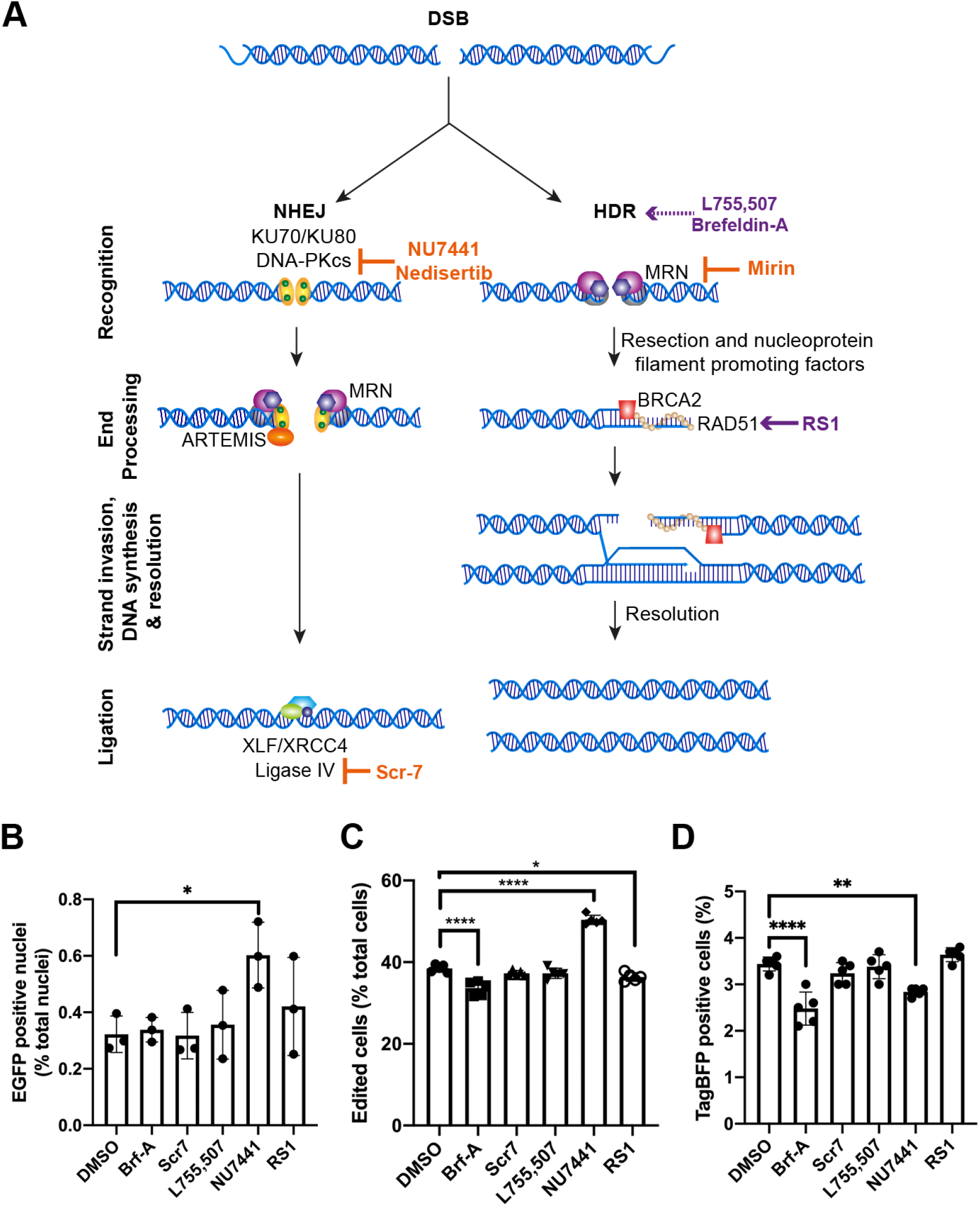
Small molecule modulators of genome editing outcome. FIVER MEFs were treated with small molecules for 24 hours post transfection: Brf-A (0.1 *μ*M), Scr7 (0.1 *μ*M), L755,507 (5 *μ*M), NU7441 (2 *μ*M) or RS1 (10 *μ*M). (A) Overview of DSB repair pathways with action of small molecules indicated. Antagonists are indicated in orange, agonists are indicated in purple. (B) EGFP positive nuclei — indicative of HDR — determined by widefield microscopy, n > 9,000 cells, N = 3 technical replicates. (C) Total observed editing, determined by flow cytometry, n = 60,000 cells, N = 5. (D) Total TagBFP^+^ cells, determined by flow cytometry, n = 60,000 cells, N = 5. Significance was tested using one-way ANOVA and Dunnett’s multiple comparisons, 0.0021 < *p* < 0.05 = *, 0.0002 < *p* < 0.0021 = **, 0.0001 < *p* < 0.0002 = ***, *p* < 0.0001 = ****. **Figure 4–Figure supplement 1. Small molecule modulators of genome editing outcome** **Figure 4–Figure supplement 2. Small molecule modulators of genome editing outcome**

Only NU7441 had a significant effect on HDR, increasing it approximately 2-fold (*p* = 0.03, one-way ANOVA with Dunnett’s multiple comparison, N = 3, **Figure 4B**). Surprisingly, NU7441 also significantly increased overall editing (**Figure 4C**) evidenced by an increase in both tdTomato^−^/EGFP^−^ and tdTomato^−^/EGFP^+^ populations (**Figure 4–figure supplement 1A and B**). This is in contrast to the decrease seen in the proportion of TagBFP^+^ positive cells after NU7441 treatment compared with DMSO control (**Figure 4D**), indicative of a reduction in NHEJ-dependent HITI. These results were recapitulated with another DNA-PKcs inhibitor (Nedisertib), which had been shown to be more effective than NU7441 (37). While Nedisertib did increase HDR (**Figure 4–figure supplement 1D**), the increase in HDR was the same as with NU7441 despite increasing total editing, tdTomato^−^/EGFP^+^, and tdTomato^−^/EGFP^−^ populations, whilst decreasing TagBFP^+^ and tdTomato^+^/EGFP^+^ populations all to a greater extent (**Figure 4–figure supplement 1E-I**). This demonstrates the ability of FIVER to rapidly and unbiasedly screen for such modulators of DNA editing outcomes.

### Rapid preclinical screening of delivery methods *in vitro*

Balancing efficacy with safety for delivery tools will be an essential part of the development of a therapeutic somatic genome editing pipeline. This requires use of relevant organotypic and pre-clinical animal models. Accordingly, FIVER was established with the aim of being a modular toolbox for streamlining the development of pre-clinical genome editing therapies for use in any relevant tissue type. Given our interest in genetic diseases of the airways, we derived FIVER primary mouse tracheal epithelial cells (mTECs), from adult reporter mice. These form stratified epithelial sheets composed of 7 cell populations (38), recapitulating the cellular environment *in vivo*. We delivered CRISPR machinery and repair constructs to mTECs varying only the method of introduction to cells using a variety of viral and non-viral lipid nanoparticle (NP) vehicles. As these cultures are representative of the *in vivo* respiratory environment, they are a powerful *ex vivo* model to prioritise respiratory epithelium tropic viral constructs or NP formulations.

We transfected FIVER mTECs using different NP formulations, composed of various lipid and peptide mixtures (39,40). These NP were used to deliver SpCas9-RNPs and MC.HDR to mTEC cultures after expansion of the basal cell population. Following maturation, mTECs were analysed for evidence of editing. For all NP formulations tested, NHEJ-based editing was observed — as both mEGFP and a loss of all membrane fluorescence (**Figure 5A**, and arrowhead). However, levels of editing were generally low and no observable HDR events were detected for any NP formulation tested (**Figure 5 – figure supplement 1A**).

**Figure 5.**
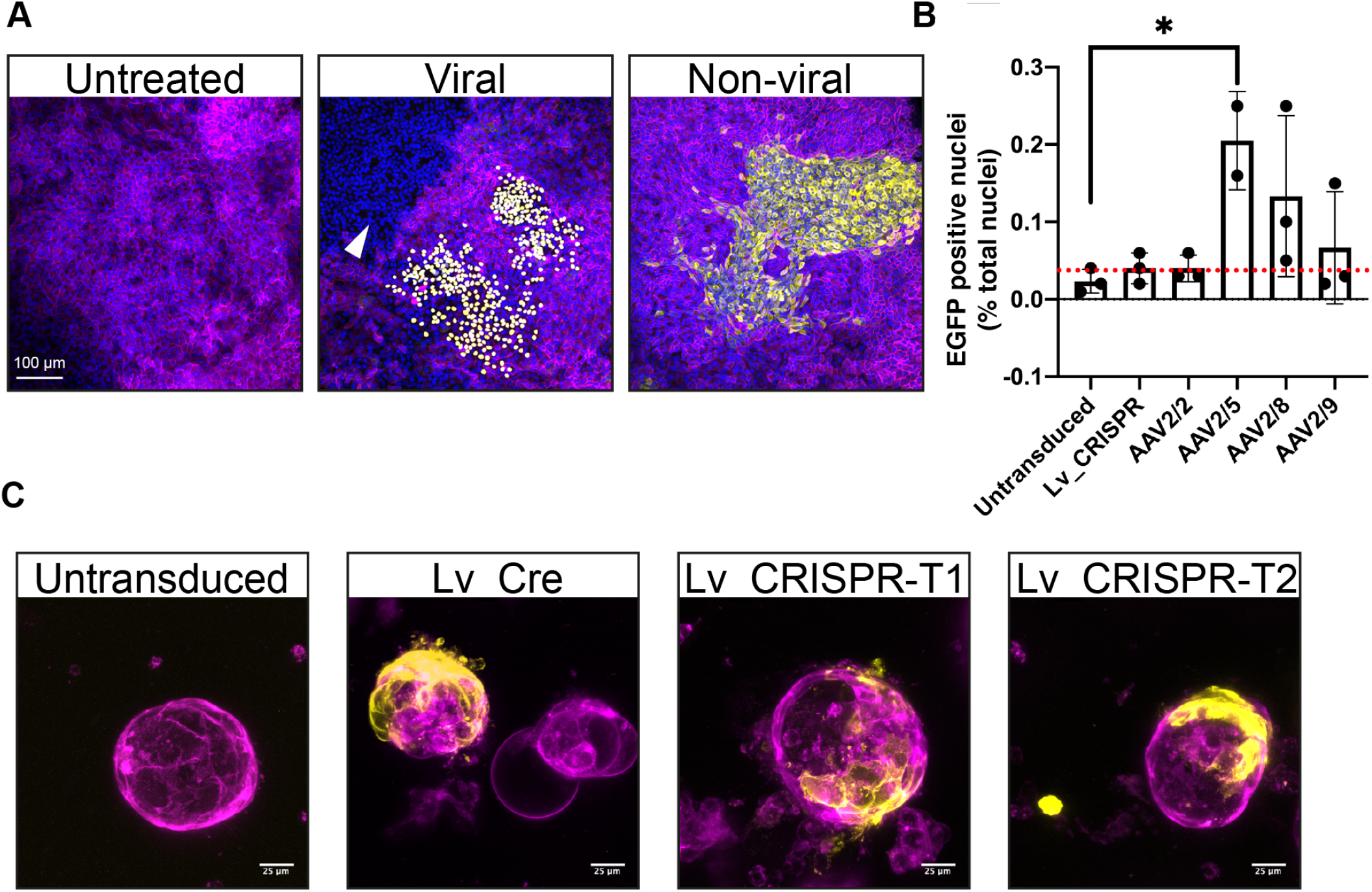
FIVER allows establishment of disease-relevant primary cultures and organoids. (A) Representative confocal images comparing viral and non-viral delivery to FIVER mTECs. Maximum intensity projections of z-stacks. For viral delivery, LV-CRISPR-T1 was combined with AAV2/5-HDR. For non-viral delivery, mTECs treated with lipid nanoparticles (DHDTMA:DOPE with peptide E) containing SpCas9/T1 RNPs and MC.HDR. NHEJ editing indicated by mEGFP fluorescence or loss of mtdTomato while HDR is illustrated by nEGFP. Arrowhead indicates tdTomato^−^/EGFP^−^ cells, also indicative of NHEJ editing. Nuclei are stained with DAPI. (B) Quantification of HDR editing in mTECs following viral transduction, n > 20,000 cells, N = 3 technical replicates, * *p* = 0.0239, one-way ANOVA with Dunnett’s multiple comparisons. Red dashed line indicates background level of detection. (C) Example confocal images of editing in FIVER ductal liver organoids. Similar activities are observed between Cre- and SpCas9-gRNA-treated organoids. Maximum intensity projections of z-stacks. **Figure 5–Figure supplement 1. FIVER allows establishment of disease-relevant primary cultures and organoids**

In parallel, we transduced FIVER mTECs with SpCas9, gRNA and an HDR template using a dual viral system. Here, the CRISPR machinery was delivered via lentivirus (with its larger packaging capacity) while the HDR templates were delivered via AAV, as AAV is particularly recombinogenic (41,42). We focused on AAV serotypes previously reported to be efficacious in delivering to airway cells (43–46) in order to determine the most efficient type for genome editing applications (**Figure 5–figure supplement 1B**). Analysis of transduced mTECs showed that all AAV serotypes tested were able to drive observable NHEJ and HDR (**Figure 5–figure supplement 1B**), though serotypes 5, 8 and 9 resulted in the greatest levels of HDR (**Figure 5B and figure supplement 1B**), while AAV2 failed to drive HDR levels above background (**Figure 5B**, dashed red line). Importantly, we were able to compare levels of editing as well as types of outcomes between viral and non-viral delivery of identical reagents, emphasising the importance of how genome editing tools are introduced into specific cell types.

Another organotypic model of translational interest is the 3D liver organoid, which allows us to bridge the gap between 2D cell cultures *in vitro* and *in vivo* studies. Self-renewing liver organoids are useful tools for disease modelling, regenerative medicine and drug screens, exhibiting genetic stability during long-term culture and some elements of liver organ physiology (47). To demonstrate the ability of FIVER to report editing in organoids, we derived 3D hepatic ductal organoids from adult FIVER mice and transduced them using lentiviruses encoding either Cre-recombinase as a positive control or CRISPR machinery. Excision of the tdTomato cassette was observed in organoids treated with either Cre or CRISPR mixes (**Figure 5C**).

### Highly eficient templated repair in FIVER early embryos

HDR is often more efficient in early embryos than in somatic cells (48,49). Thus, to demonstrate our reporter in an optimal system, we investigated the amount and type of genome editing outcomes in blastocysts following nuclear microinjection of FIVER single cell zygotes; we carried out pronuclear injections using SpCas9-RNPs and minicircle repair templates (MC.HDR or MC.HITI). Embryos were cultured for 72 hours onto blastocyst stage where confocal imaging revealed high levels of all editing events (**Figure 6A and B**).

**Figure 6.**
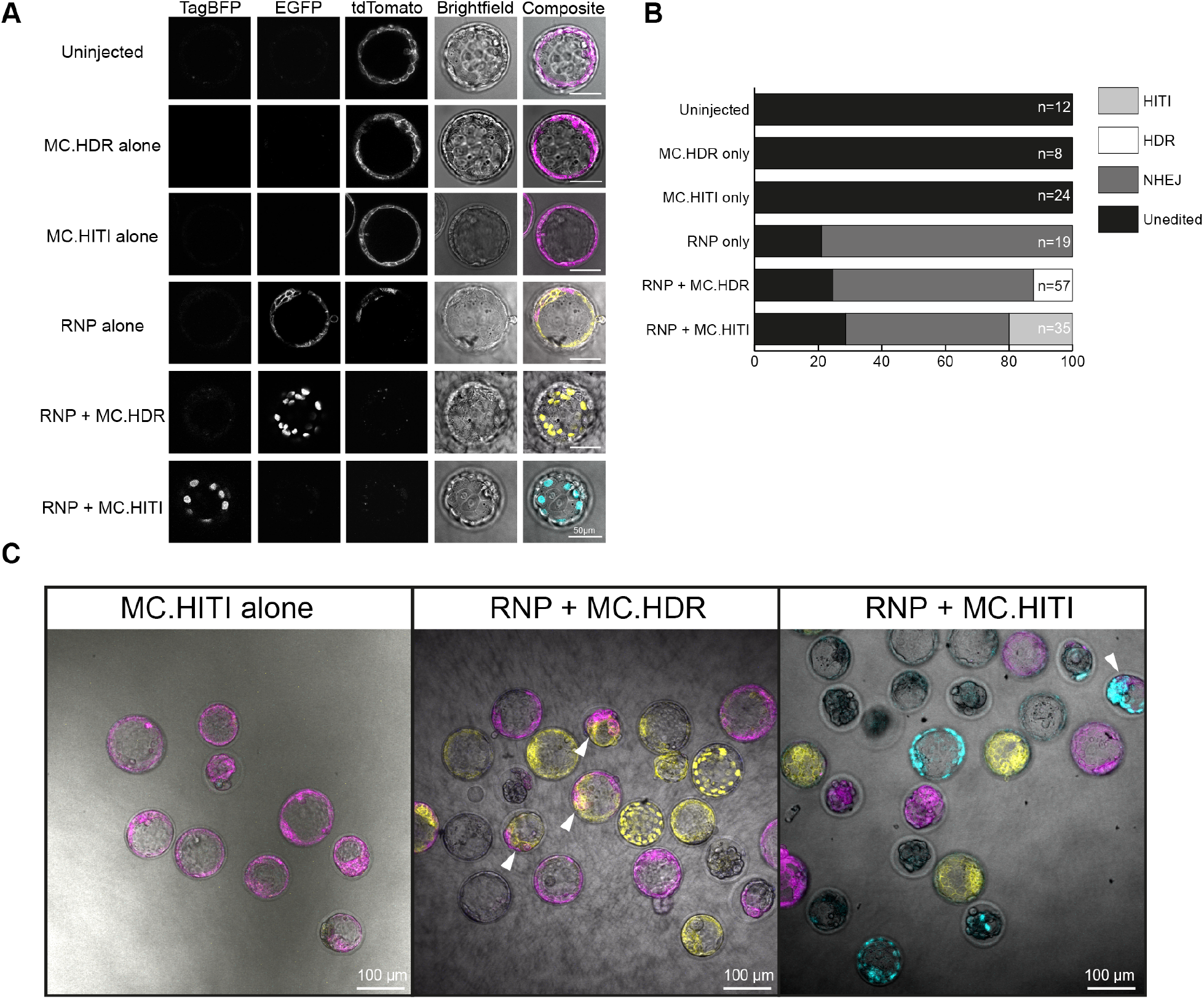
Highly eficient editing in FIVER early embryos. SpCas9-T1 RNP and minicircle repair constructs (MC.HDR or MC.HITI) were delivered to FIVER single cell zygotes by pronuclear injections. After progression to blastocysts (72 hours post single cell injection), they were analysed by confocal microscopy. (A) Representative confocal images indicating the ability of FIVER to demonstrate all editing outcomes. Scale bars represent 50 *μ*m. Single z-slices are presented. (B) Quantification of all editing outcomes. Total numbers of blastocysts in each group are indicated, from 2 rounds of injections. Blastocysts that arrested were discounted from analysis. (C) Representative confocal images of edited blastocysts indicating the range of editing outcomes observed. Arrowheads indicate mosaic editing events. Single z-slices. For full z-slice montage, see **Figure 6–video 1**. **Figure 6–video 1. Single slice montage of eficient editing in FIVER early embryos**. Videos show full z-slice montage of confocal images through blastocysts, cultured 72 hours post single cell injection. Field of blastocysts following injection with (A) HITI only, (B) RNP only, (C) RNP + MC.HDR or (D) RNP + MC.HITI. Scale bar 100*μ*m. See also **Figure 6**.

In the majority of cases (88/110, 80%), blastocysts demonstrated editing in all cells using RNPs (**Figure 6**). In a small subset (22/110, 20%), mosaic editing was observed (**Figure 6C**, arrowheads and **Figure 6–video 1**), indicative of a delay in the initial editing event past the one cell stage. In early embryos, the rates of HDR and HITI were similar, compared to asynchronous primary FIVER fibroblasts cultures where HITI was 10-fold more efficient than HDR (**Figure 6B** versus **Figure 4B and D**). This demonstrates that by using the same reagents in different cells types, FIVER can track how different cell types differ in their predominant choice of repair mechanism.

### Tracking genome editing events *in vivo* following hydrodynamic tail vein injection

The major advantage of our FIVER model is the ability to monitor *in vivo* editing spatially and temporally in any tissue of interest. To capitalise on this, we delivered CRISPR based editing machinery and repair constructs to adult mice via hydrodynamic tail vein injection (HTVI) using naked DNA constructs (**Figure 7A**) (50). HTVI involves a rapid injection of a large volume into the animal causing a transient disruption of the microvascular barrier in the liver sinusoids such that DNA is rapidly absorbed by hepatocytes. We inserted our editing machinery into a plasmid-based *Sleeping Beauty* (SB) transposon vector (SB-CRISPR) which is able to efficiently integrate its transgene cargo into the genome of targeted cells (51). The SB transposon utilizes a random integrative cut-and-paste transposition mechanism, where its integration site profile is not biased towards actively transcribing genes unlike lentiviral vectors (52–54). Livers were harvested 1 week post injection and analysed for evidence of editing using confocal microscopy.

**Figure 7.**
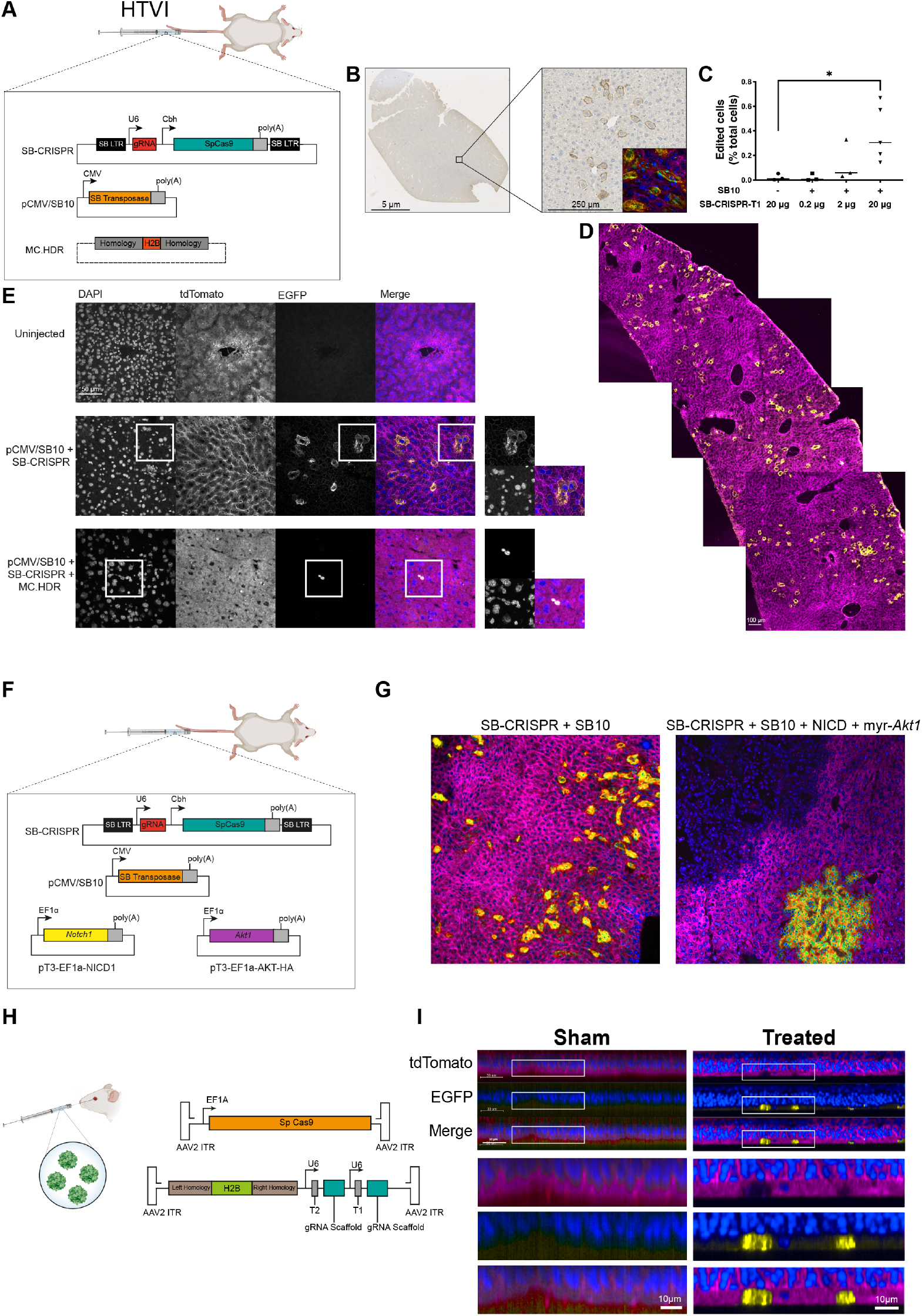
FIVER reports on *in vivo* editing in multiple organ systems. (A) Plasmid and minicircle constructs used for HTVI. (B) Wax sections of liver stained with anti-GFP antibodies, used to quantify overall levels of editing following administration of varying amounts of SB-CRISPR-T1. (C) Quantification of total editing (EGFP positive cells/ total cells). The presence of SB10 transposase significantly increases the level of editing with 20 *μ*g SB-CRISPR-T1. * *p* = 0.0329, one-way ANOVA with Dunnett’s multiple comparisons. (D) Composite maximum intensity projection of a confocal image, illustrating widespread liver editing following HTVI. (E) Representative confocal images of liver sections from HTVI mice. Magnified sections indicate NHEJ (mEGFP) and HDR (nEGFP) editing outcomes. Maximum intensity projection of z-stacks. (F) Overview of constructs used in the HTVI liver tumour model. (G) Representative confocal images to show liver tumour development. Tumours display editing in the FIVER mice, by either gaining mEGFP or losing mtdTomato fluorescence. No tumours observed in control animals not injected with NICD and myr-*Akt1*. Nuclei are stained with DAPI. NICD = *Notch1* intracellular domain, myr-*Akt1* = myristoylated *Akt1*. Maximum intensity projection of z-stacks. (H) Overview of viral constructs delivered subretinally to FIVER mice. (I) Representative confocal microscopy of retinal wholemounts following subretinal delivery of AAV. Eyes harvested 14 days after injection into post-natal day 3 animals. Nuclei stained with DRAQ5 are indicated in blue.

By using different amounts of the SB-CRISPR-T1 plasmid, we demonstrate that there is a correlation between the amount of the CRISPR machinery delivered and the level of editing observed (**Figure 7B-C**). Editing is only observed when the SB10 transposase (55) is also present. Consequently, we found that 20 *μ*g of SB-CRISPR-T1 was optimal and this amount was used in all subsequent experiments.

In sham treated animals, there was no evidence of editing — all liver sections analysed retained mtdTomato fluorescence (**Figure 7E**). In all SpCas9-gRNA treated cases, NHEJ editing was evident throughout the postnatal livers — indicated by the switch from mtdTomato to mEGFP (**Figure 7D-E**). In addition, nEGFP expression was observed in animals that received HDR repair templates, indicating that low levels of HDR had occurred (**Figure 7E**). Given that the bulk of adult hepatocytes are post-mitotic, low levels of HDR are predicted, but injury from the HTVI could possibly trigger cell cycle re-entry.

### FIVER facilitates tracking the fate of edited cells *in vivo*

Another important application of genome editing has been to screen *in vivo* for genetic drivers of tumorigenesis in mouse models (51). Given the complexity of delivering a library of gRNAs and nucleases to many different cell types and tracking their fates over time, we postulated that FIVER could aid in such screens by allowing lineage tracing of edited cells. Following co-delivery of a library of genome-wide gRNAs along with T1 gRNA, we aimed to track edited tumours by a shift in fluorescence. Hits which increased or decreased tumour pathology, marked by a change in fluorescence, would be of interest for further study. This would enable isolation of mutant cells prior to establishment of frank carcinoma and also allow for more in-depth analyses of tumour progression, as opposed to current methods which examine loss of function mutations solely in established tumours. We therefore co-delivered the SB-CRISPR-T1 and pCMV/SB10 plasmids with two drivers of tumorigenesis — *Notch1* receptor intracellular domain (NICD) and *Akt1* containing a myristoylation sequence (myr-*Akt1*) (56), via HTVI (**Figure 7F**). After 6 weeks, livers were analysed for evidence of tumours showing a shift in fluorescence.

In all cases, tumours were observed only when the oncogenes were provided (**Figure 7G**). When analysed by confocal microscopy, tumours were shown to be either tdTomato^−^/EGFP^+^ or lacking in all fluorescence (tdTomato^−^/EGFP^−^), both outcomes indicative of NHEJ editing (**Figure 7G**). Changes in fluorescence upon editing will greatly aid in resecting tumour cells out from non-edited stroma for clean genotyping and expression profiling.

### Eficient retinal editing following subretinal AAV administration

Given its accessibility and compartmentalisation, the eye represents a leading target tissue for gene therapies, including genome editing (57,58). To demonstrate the ability of FIVER to accelerate the development of such therapeutic approaches, we carried out subretinal injections of AAV-based CRISPR machinery in neonatal FIVER mice (**Figure 7H**).

Following injection, animals were allowed to recover for 14 days, then sacrificed and eyes analysed for editing. All mice treated with AAVs demonstrated retinal NHEJ editing — transition from mtdTomato to mEGFP (treated, **Figure 7I**) — while sham injected animals retained mtdTomato fluorescence throughout (sham, **Figure 7I**).

### Editing outcomes at the FIVER reporter locus faithfully relect editing outcomes at a second independent locus

Visualisation of genome editing outcomes across tissues and whole organisms will help expedite development of more efficient and better tolerated delivery systems for somatic genome editing tools and more efficacious therapeutics. However, the question remains whether editing outcomes at the FIVER locus — *Rosa26*, which is ubiquitously expressed in mouse — would be indicative of what happens at a second locus of therapeutic interest, that may not be widely expressed. Chromatin accessibility and modifications have been reported to have variable effects on the efficacy and type of editing outcomes (59–63). To address this, we crossed the FIVER mice with a preclinical model of primary ciliary dyskinesia (PCD), harbouring a 7-bp deletion in the *Zmynd10* gene (*Zmynd10^em^*^1^*^Pmi^*) (64). From these mice, we generated *FIVER/Zmynd10^em^*^1^*^Pmi^*) MEFs which we transfected with SpCas9-RNPs targeting both FIVER and *Zmynd10* with corresponding MC.HITI repair constructs (**Figure 8B**).

**Figure 8.**
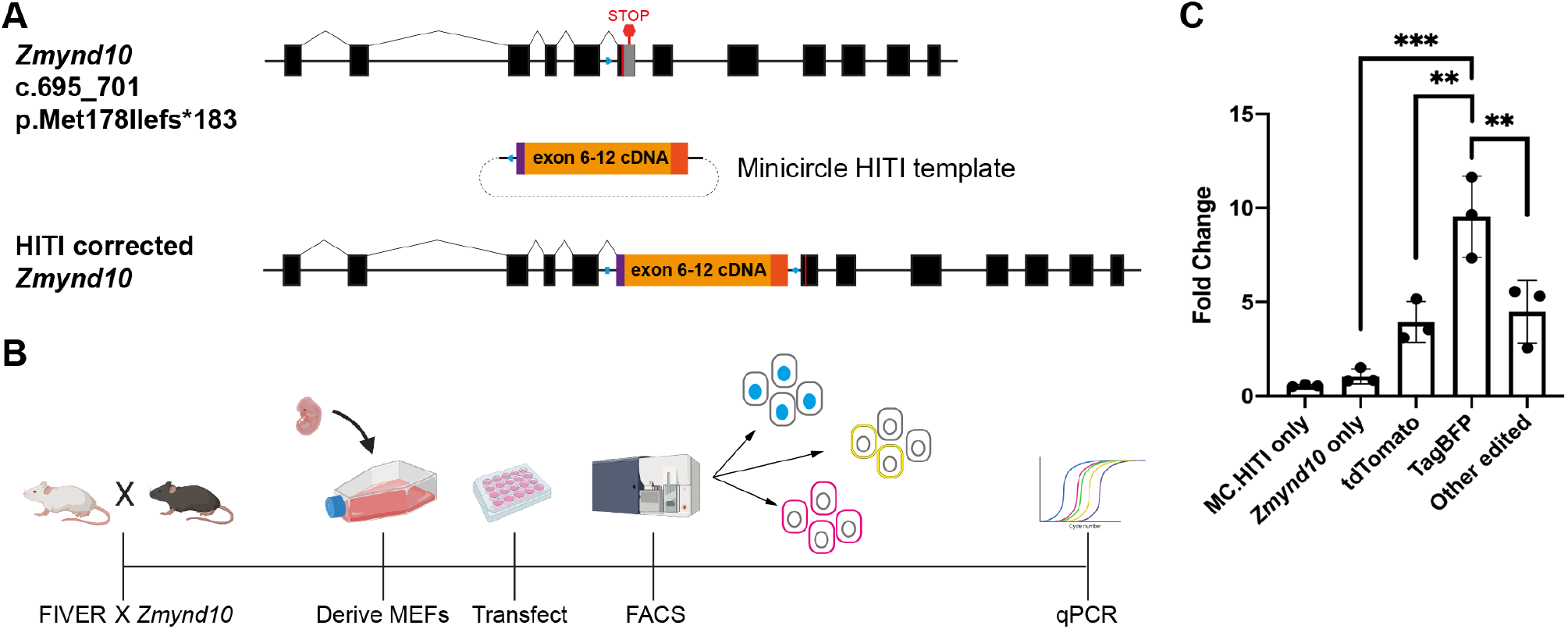
Editing outcomes at FIVER recapitulate editing at a second independent disease locus, the ciliopathy gene *Zmynd10*. (A) Overview of expected HITI editing outcome at *Zmynd10*. Blue polygon depicts gRNA target site in both the target locus and the repair construct. Upon correct integration the gRNA site is destroyed, leaving two remnant sites (blue rectangle and blue diamond), which can no longer be recognised by the gRNA. (B) Experimental workflow. (C) Overall HITI editing at *Zmynd10* locus in sorted or *Zmynd10*-targeted alone populations. One-way ANOVA with Tukey’s multiple comparisons, N = 3 technical replicates, ** *p* = 0.0029 (TagBFP vs. tdTomato) or *p* = 0.006 (TagBFP vs. Other edited), *** *p* = 0.0001.

Cells were sorted into three populations by FACS: tdTomato single positive (tdTomato^+^/EGFP^−^; 72.2%), TagBFP single positive (tdTomato^−^/EGFP^−^/TagBFP^+^; 3.1%), and the third population consisting of all other fluorescent outcomes (tdTomato^−^/EGFP^+^, tdTomato^−^/EGFP^−^ and tdTomato^+^/EGFP^+^; 16.1%) (**Figure 8C**). In addition, a population treated with only *Zmynd10*-targeting SpCas9-RNP and MC.HITI was included. While lower overall levels of editing were observed here — 16.1% total edited cells and 3.1% TagBFP^+^ cells — versus previous experiments (**Figure 4C and D**), these cells were supplied with a 50% lower concentration of editing reagents targeting FIVER and hence this would be expected given the correlation between reagent dose and levels of editing (**Figure 7C**). qPCR to detect integration of the HITI cassette at *Zmynd10* revealed a significant (*p* = 0.006, one-way ANOVA with Tukey’s multiple comparison) 10-fold enrichment of HITI editing at the *Zmynd10* locus within the TagBFP single positive population when compared to all other populations (**Figure 8C**). These results suggest that FIVER should be a powerful, widely-applicable tool to track specific editing outcomes at different loci in different cell populations *in vivo*.

## Discussion

Here, we have developed and characterised a novel, multispectral fluorescent reporter of *in vivo* genome editing — FIVER. We believe it to be the first of its kind to sensitively report editing outcomes *in vitro* and *in vivo* for NHEJ, HDR and HITI editing outcomes. We confirm by deep sequencing that changes in fluorescence emission and/or localisation broadly and faithfully recapitulate underlying genomic changes. These changes at the genomic level result in rapid and biphasic changes in fluorescence, which are fully complete within 48 hours for all observed outcomes (**Figure 3**). We show that FIVER’s fluorescent read-out quantifiably reflects changes at the DNA level in multiple primary cell types and complex tissues.

The field of genome editing is rapidly evolving with new and improved nuclease tools; a broader genomic range we can target with different PAM sites, improved specificity, and novel mechanisms of action and resolution of DNA breaks (65,66). FIVER can also be used with other genome editing platforms including TALENs (for a list of potential TALEN target sites see **Supplementary Table 1**) and other Cas proteins, as long as they introduce DSBs. Different nucleases leave different ends at DSBs and how these are resolved may bias the outcomes. For example, Cas9 generates predominantly blunt ends, whilst Cas12a generates sticky ends (67,68); the latter are suggested to be more amenable to targeted knock-in strategies. The FIVER toolbox can be rapidly expanded to include novel nucleases to explore their efficiencies and the editing outcomes they elicit *in vivo*, as they are taken forward for preclinical use.

Using FIVER, we investigated a range of previously reported small molecule modulators of DSB repair. In our initial screen, only NU7441 significantly increased HDR (**Figure 4B**). In addition, we also observed a significant reduction in the number of TagBFP^+^ cells, confirming that HITI results from NHEJ-mediated knock-in of the repair template (**Figure 4D**). Though counter-intuitive, NU7441 treatment also increased the level of overall editing, by increasing both tdTomato^−^/EGFP^−^ and tdTomato^−^/EGFP^+^ populations (**Figure 4B and figure supplement 1A and B**). The increase in tdTomato^−^/EGFP^+^ could be accounted for to some extent by the concomitant increase in HDR (nEGFP). However, the tdTomato^−^/EGFP^−^ population is believed to result from imprecise NHEJ repair such that we would see a reduction of this population following inhibition of DNA-PKcs. As we observed in the NGS data, this population results from larger deletions following excision of the tdTomato cassette which extend into the promoter region or coding sequence of EGFP (**Figure 2–figure supplement 1B**). Mutations such as these may be the result of alt-NHEJ, specifically microhomology-mediated end-joining (MMEJ), which is known to result in larger indels than canonical NHEJ (69). Indeed, Schep et al. have recently demonstrated that inhibition of NHEJ using NU7441 leads to an increase in the proportion of MMEJ-mediated repair (63), suggesting MMEJ may similarly compensate for a reduction in NHEJ, consistent with our results. Inhibition of MMEJ with mirin (70) had a similar, though less pronounced effect as NU7441 on both tdTomato^−^/EGFP^+^ and tdTomato^−^/EGFP^−^ populations, but in combination with NU7441 it was cytotoxic (**Figure 4–figure supplement 2**), suggesting inhibition of multiple DSB repair pathways is not tolerated. In addition, our NGS data revealed asymmetry in editing between the two gRNA targets sites, with more indels present at the upstream site (**Figure 2B**). This implies that editing at the two near identical sites could be asynchronous or that local sequence differences lead to more disruptive repair at the upstream site. Taken together, these suggest that multiple repair pathways may be employed following CRISPR activity and that blocking one or more merely shifts the balance between these competing pathways.

We also investigated Nedisertib, reported to be a more potent inhibitor of DNA-PKcs (37). However, we found that Nedisertib was less efficacious than NU7441 at increasing HDR after 24 hours of treatment, though was a more potent inhibitor of HITI (**Figure 4–figure supplement 1D and F**). Considerable controversy still exists about how DSBs elicited by genome editors are resolved and the molecular mechanisms involved. The majority of these small molecule studies have been done in cancer cell lines, with replication studies in alternative cell lines failing to recapitulate findings (65); it remains to be seen whether similar pathways are employed in primary cells. Whether cell type specific differences exist in regulators of these pathways also remains unclear. Being able to control or bias editing outcomes with small molecule modulators is attractive. FIVER would be a powerful way of verifying efficacy and toxicity of known drugs in target cells of interest, as well as offering the opportunity to screen for novel candidates in an automated fashion, taking advantage of fluorescent shifts and localisations. As part of the FIVER toolkit, we have developed automated quantification scripts in QuPath (open source) to aid with these types of applications; these are available on GitHub (https://tinyurl.com/ycbcoopk). In addition, FIVER allows testing in other relevant cells, tissues and ultimately *in vivo*.

One of the major applications for FIVER will be in optimising delivery of genome editing tools to different cell and tissue types *in vivo*. In contrast to gene augmentation studies, high level, prolonged expression of genome editing tools is likely not desirable in therapeutic settings. A short, but widespread, burst of editing activity is likely ideal to avoid off-target effects such that editing can be biased towards the desired outcomes. Preclinical studies to explore how best to balance efficacy (i.e., efficient editing) and safety (i.e., high on-target, non-integrating activity) are needed. Using FIVER, we were able to demonstrate that even identical gRNA and Cas9 nuclease complexes elicited very different outcomes in our airway organotypic cultures; with AAV-delivered HDR repair effecting robust editing and greater HDR compared to nanoparticle delivery at a proliferative stage (**Figure 5A and figure supplement 1A and B**). However, these nanoparticle reagents were optimised for targeting mature airway epithelium (40), where the bulk of cells would be differentiated and likely less amenable to HDR. FIVER will be a powerful tool to unbiasedly isolate edited cell populations following *in vivo* editing by imaging or FACS-based methods. This will allow researchers to determine which cell types have been edited, quantify at what levels and determine their distribution within the tissue (i.e., proximal to distal in the airways), plus their biodistribution in the organism. FIVER will also allow us to address important questions such as the extent to which edited cells undergo clonal expansion and how long edited cells remain in the tissue, during health and in disease models.

Crucially, we were able to demonstrate HITI editing outcomes at a second independent locus of clinical interest (**Figure 8C**). HITI editing has great potential as a therapeutic approach in many genetic diseases. Achieving therapeutic levels of perfect repair by HDR is still a substantial hurdle for the development of genome editing-based therapeutics. However, as HITI takes advantage of the more prevalent NHEJ pathway, it can help to bridge the gap between the precise editing of HDR and the variable indels generated by NHEJ, resulting in a more predictable, targeted repair which occurs more efficiently than HDR strategies. HITI is also a more realistic repair strategy in non-proliferative cell types. Indeed, in their study, Suzuki et al. were able to demonstrate potential therapeutic benefit of HITI by restoration of the *Mertk* gene in a rat model of retinitis pigmentosa (19). HITI-targeted animals showed greater improvements in retinal morphology and in both rod and cone function compared to HDR-targeted animals. More recently, others have shown the potential of HITI for use with other Cas proteins; an AAV based HITI strategy making use of SaCas9 was shown to restore FIX serum levels to a greater extent than the equivalent HDR strategy in a mouse model of haemophilia (71). Furthermore, HITI can be used to aid in gene augmentation therapies — targeting genes to safe harbour loci for sustained expression without the risk of insertional mutagenesis (72). The ability of FIVER to report HITI editing will be beneficial in developing new and improved HITI-based therapeutics.

There are currently several fluorescence-based editing reporters available, however the majority of these are limited to *in vitro* use (10–14,16,17). While a few *in vivo* editing reporters have also been described, these are limited to reporting on NHEJ outcomes (73,74). Whilst this work was in preparation, Alapati et al. reported using the *mTmG* reporter to monitor NHEJ editing outcomes — solely *in utero* — with adenovirus delivery for a rare genetic lung disease (15). We believe repurposing this readily available fluorescent reporter system for genome editing with the robust FIVER toolbox to report on NHEJ, HDR and HITI outcomes *in vivo* creates a valuable community resource which will expedite effective genome therapies. In addition, the availability of well-established preclinical mouse models of human disease enables rapid introduction of the reporter into physiologically or pathologically relevant animals. As such, FIVER has the potential to accelerate the development of effective genome surgery across a broader spectrum of genetic diseases.

FIVER will allow vectors, vehicles and small molecule modulators to be tested by independent labs, and evolving methods and reagents that improve outcomes following ‘genome surgery’ can be shared for everyone’s benefit.

## Methods and Materials

### Plasmids

The following plasmids were a gift from Feng Zhang: pX330, (Addgene plasmid #42230; http://n2t.-net/addgene:42230; RRID: Addgene_42230); pLentiCRISPRv2, (Addgene plasmid #52961; http://n2t.-net/addgene:52961; RRID: Addgene_52961) and pAsCpf1(TYCV)(BB) (pY211), (Addgene plasmid #89-352; http://n2t.net/addgene:89352; RRID: Addgene_89352). The piRFP670-N1 plasmid was a gift from Vladislav Verkhusha (Addgene plasmid #45457; http://n2t.net/addgene:45457; RRID: Addgene_45457). The SB-CRISPR plasmid was a gift from Ronald Rad. The pCMV/SB10 plasmid was a gift from Perry Hackett (Addgene plasmid #2455; http://n2t.net/addgene:24551; RRID: Addgene_24-551). Both pT3-myr-AKT-HA (Addgene plasmid #31789; http://n2t.net/addgene:31789; RRID: Addgene_31789) and pT3-EF1a-NICD1 (Addgene plasmid #46047; http://n2t.net/addgene:46047; RRID: Addgene_46047) were gifts from Xin Chen. Oligonucleotides containing the gRNA sequences were synthesised by Sigma-Aldrich (USA) (Table 2) and cloned into pX330, SB-CRISPR or pAsCpf1(TYCV)(BB) (pY211) following digestion with BbsI restriction endonuclease. pLentiCRISPRv2 was engineered to contain the iRFP670 fluorescent protein downstream of Cas9 using a self-cleaving peptide motif (P2A). The same gRNAs were cloned into pLentiCRISPRv2-iRFP670 following digestion with BsmBI. All gRNA sequences are detailed in **Table 1**.

**Table 1.**
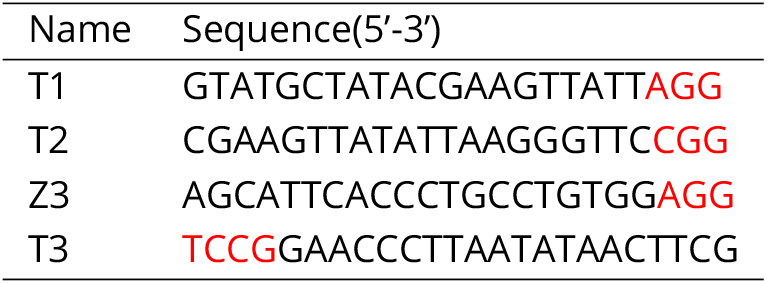
gRNA Sequences. Target sequences are given in black, with PAMs given in red.

**Table 2.**
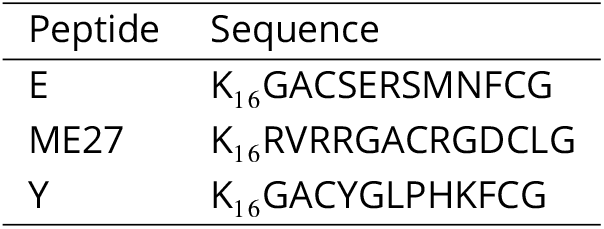
Peptides used for lipid nanoparticle formulation. Peptides E and Y are epithelial targeting peptides (40) and ME27 is an RGD-containing integrin-targeting peptide.

| Peptide | Sequence |
| --- | --- |
| E | K<sub>16</sub>GACSERSMNFCG |
| ME27 | K<sub>16</sub>RVRRGACRGDCLG |
| Y | K<sub>16</sub>GACYGLPHKFCG |

### Viral vectors

AAV vectors were produced by Virovek (USA). Lentiviral vectors, all coated with VSV-G, were produced by the Viral Vectors Core at the Shared University Research Facilities, the University of Edinburgh (Edinburgh, UK).

### Minicircle DNA vectors

Production of minicircle vectors was carried out by PlasmidFactory (Germany). Sequences are listed in **Supplementary sequences**.

### Cell culture

Mouse embryonic fibroblasts (MEFs) were derived from embryonic day 11.5 to 13.5 (E11.5 - E13.5) FIVER embryos. Cells were cultured in Opti-MEM supplemented with 10% v/v foetal calf serum and 1% v/v penicillin/streptomycin, at 37°C, 5% CO_2_ in a humidified incubator. For immortalisation, these were transfected with a plasmid containing SV40 large T antigen and selected for using puromycin (3 *μ*g/mL).

Mouse tracheal epithelial cells (mTECs) were derived from tracheas of 5-7 week old FIVER mice. Basal cell populations were first expanded in KSFM media (Gibco, USA) supplemented with 1% v/v penicillin/streptomycin, 0.025 *μ*g/mL murine epidermal growth factor (Scientific Laboratory Supplies, UK), 0.03 mg/mL bovine pituitary extract (Gibco, USA), 1 *μ*M isoproterenol (Sigma-Aldrich, USA), 10 *μ*M Y-27632 (StemCell Technologies, UK) and 5 *μ*M DAPT (Sigma-Aldrich, USA). Cells were then cultured on semipermeable supported membranes (Transwell; Costar, USA), as previously described (75). 10 *μ*M Y-27632 (StemCell Technologies, UK) was added to the medium during the proliferation stage to promote basal cell proliferation.

### Organoid culture

Hepatic organoids were generated from isolated bile ducts. Briefly, isolated bile ducts from outbred adult FIVER mice were resuspended in 100% GFR Matrigel, plated in base media consisting of DMEM/F-12 supplemented with Glutamax, Penicillin/Streptomycin, Fungizone and HEPES (ThermoFisher Scientific, USA). These were allowed to expand at 37°C, 5% CO_2_ in a humidified incubator. Following expansion, ducts were removed from Matrigel by incubating with ice-cold Versene and dissociated with pipetting, before re-plating in fresh 100% Matrigel. This process was repeated to expand organoids. Just prior to feeding, the base media was supplemented with HGF, EGF, FGF10, Gastrin, Nicotinamide, N-Acetylcystine, B-27, Forskolin, Y-27632 (StemCell Technologies, UK), A83-01 (TGF-*β* inhibitor) and Chir99021 (GSK3*β* inhibitor).

### Transfections and Transductions

All nucleofections were carried out using the Neon transfection system (ThermoFisher Scientific, USA). For small scale plasmid transfections, 10 *μ*L tips were used. A total of 1 *μ*g DNA and 0.5 ×10^5^ cells were transfected per tip using 1350V, 30ms and a single pulse. For large scale plasmid transfections, 100 *μ*L tips were used with 10 *μ*g DNA and 1 ×10^6^ cells per tip. Transfection using RNPs were carried out using the same Neon conditions, using a total of 1 *μ*g of Cas9 protein (ThermoFisher Scientific, USA) per 0.5 ×10^5^ cells.

Ribonucleoprotein complexes (RNPs) were generated using GeneArt Platinum Cas9 nuclease (ThermoFisher Scientific, USA) and in vitro transcribed gRNA in a ratio of 1 *μ*g Cas9:240 ng gRNA. Complexes were allowed to form at room temperature for 5-10 min prior to use. gRNA was produced using the GeneArt Precision gRNA synthesis kit (ThermoFisher Scientific, USA), according to the manufacturer’s instructions.

For lipid nanoparticle-based transfections, nanoparticles were generated using a weight ratio of 1:1:4 (Cargo:Lipid:Peptide, where cargo is RNP complexes with or without MC.HDR). The lipid component was either 2,3-dioleyloxypropyl-1-trimethyl ammonium chloride (DOTMA) or 1-propanaminium, N,N,N-trimethyl-2,3-bis (11Z-hexadecenyloxy)-iodide (DHDTMA), mixed 1:1 in a molar ratio with the neutral lipid dioleoyl L-*α*-phosphatidylethanolamine (DOPE) (39). The peptides used are listed in **Table 2**. Complexes were allowed to form for 30 min at room temperature, diluted in OptiMEM and applied to cells. Plates were centrifuged at 1500g for 5 min. Cells were incubated at 37°C, 5% CO_2_ in a humidified incubator for 4 hours before complexes were removed and fresh media applied.

For viral transduction of mTECs, 10 *μ*L of each virus was diluted in growth media containing polybrene (10 *μ*g/mL; Sigma-Aldrich, USA) then mixed with cells, incubated at room temperature for 10 min then plated onto transwell membranes as described above. Lentivirus was added at 1.5 × 10^11^ TU/mL and AAVs were used at 1 × 10^13^ vg/mL. For transduction of hepatic organoids, lentivirus was diluted in base media containing polybrene (10 *μ*g/mL; Sigma-Aldrich, USA) and added directly to organoid cultures.

### Small molecule treatments

The following small molecule modulators of genome editing outcome were used in this study: Brefeldin A, L-755,507, NU7441, M3814 (Nedisertib), RS-1 and mirin (B012-5mg, 18629-5 mg-CAY, 14881-5 mg-CAY, HY-101570-10mg, B1118-5 and 13208-5 mg-CAY, respectively; Cambridge BioScience, UK), and SCR7 (M60082-2s, XcessBio, USA). All were reconstituted in DMSO. For use in tissue culture, each drug was diluted to a final working concentration (as indicated) alongside a DMSO only control and added to cells immediately after transfection for a period of 24 hours.

### Fluorescence activated cell sorting

Cells were detached using TrypLE Express reagent (ThermoFisher Scientific, USA), pelleted by centrifugation and resuspended in PBS. For analysis, a BD LSRFortessa was employed, for sorting, either a BD FACSJazz or BD FACSAria were used (all BD Biosciences, USA). For EGFP an excitation filter of 488/50 was used with an emission filter of 525/50 (488-525/50). For tdTomato, 561-610/20, 561-586/15 or 561-582/15 were used depending on the machine. For TagBFP 405-450/50 was used. For analysis, a total of 50-100,000 cells were used.

### Sequencing

DNA was extracted from cells using the DNeasy Blood and Tissue Kit (QIAGEN), according to manufacturer’s instructions. For NGS, the primers are listed in **Table 3**. Sample preparation and sequencing was carried out by Edinburgh Wellcome Trust Clinical Research Facility (WTCRF). Briefly, amplicons were quantified using a Qubit dsDNA HS kit or BR assay (Ion Torrent and MinION, respectively; ThermoFisher Scientific, USA). For Ion Torrent, these were sheared using a Covaris E220 Evolution Focused Ultrasonicator (ThermoFisher Scientific, USA), quantified and barcoded. The library was then amplified (10 cycles) and size selected using AMPure XP beads (Beckman Coulter, California, US) for fragments approximately 300bp in length, checked for purity, quantified, and an equimolar stock was prepared and sequenced.

**Table 3.**
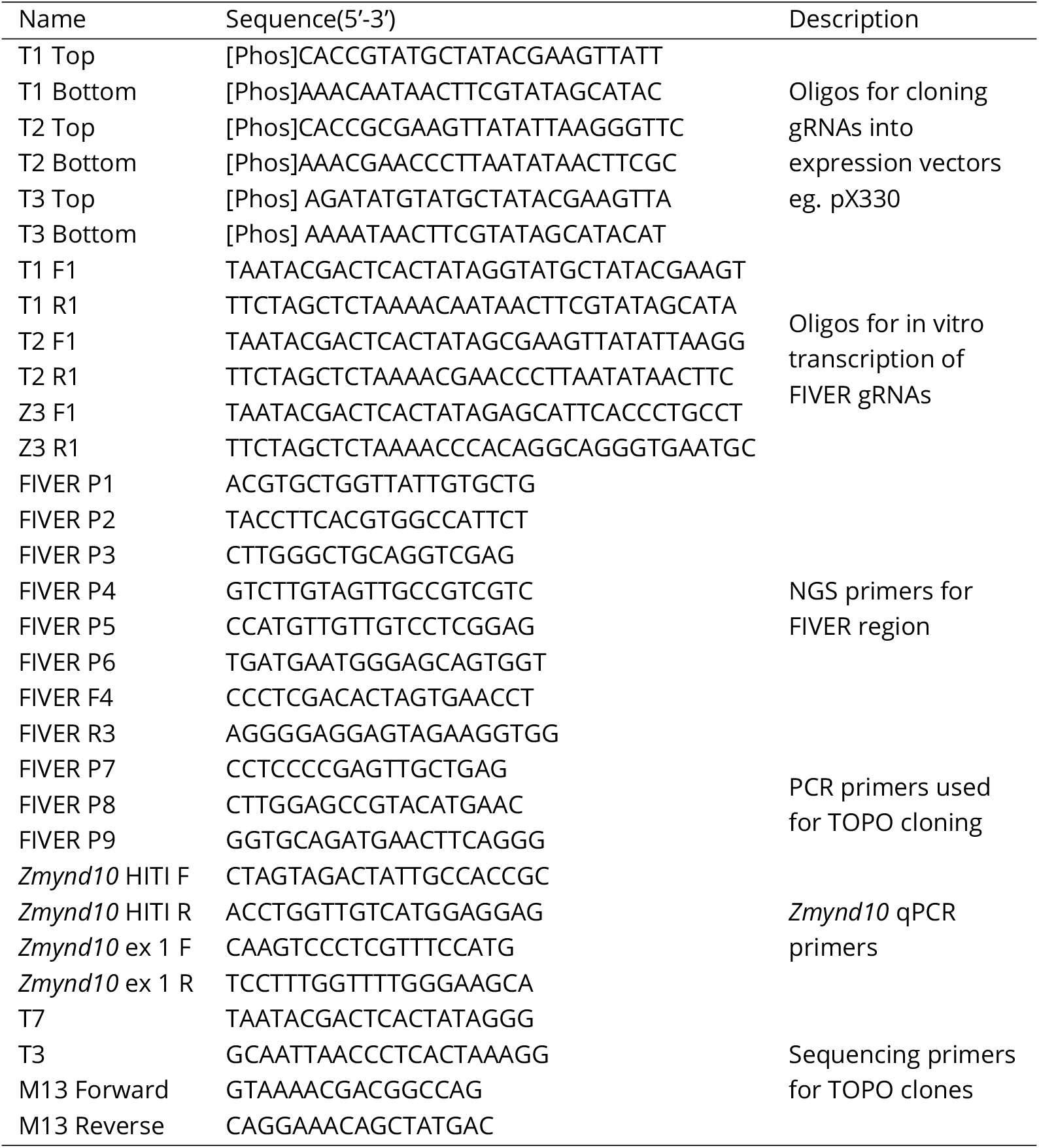
Oligonucleotide sequences

For MinION, 50 ng of each amplicon was end-repaired and adenylated using an NEBNext Ultra End Repair/dA-Tailing Module kit (NEB, USA) and purified using AMPpure XP beads (Beckman Coulter, USA). Barcode adapters from the PCR Barcoding Kit 96 (Oxford Nanopore Technologies, UK) were ligated to the end-repaired, dA-tailed DNA during 18 cycles of PCR. Excess barcode adapters were removed using AMPure XP beads, and barcoded DNA was quantified using a Qubit dsDNA HS assay (ThermoFisher Scientific, USA). Equal quantities of each barcoded amplicon were pooled before being end-repaired and adenylated to allow ligation of sequencing adapters and tethers from the Nanopore 1D2 Sequencing Kit (Oxford Nanopore Technologies, UK). Libraries were re-purified and an equimolar stock was prepared and sequenced.

For targeted sequencing of HDR samples, PCR amplification of the whole locus was carried out using the following primers: FIVER F4 and FIVER R3 (**Table 3**). Products were purified using the PureLink quick PCR purification kit (ThermoFisher Scientific, USA) according to the manufacturer’s instructions. 4 *μ*L of purified product was cloned into the pCR-4 Blunt TOPO vector using the Zero Blunt TOPO PCR Cloning Kit for Sequencing (ThermoFisher Scientific, USA). To identify larger deletions in the promoter region, PCR amplifications using P7 and P8 or P7 and P9 primers was carried out (**Table 3**). Products were purified using the PureLink quick PCR purification kit (ThermoFisher Scientific, USA) according to the manufacturer’s instructions. 4 *μ*L of purified product was cloned into the pCR-4TOPO vector using the TOPO TA Cloning Kit for Sequencing (ThermoFisher Scientific, USA). In both cases, colonies were selected and grown overnight at 37°C, 300 rpm in 96-well plate cultures in LB containing 100 *μ*g/mL ampicillin. DNA extraction and sequencing were performed by the IGMM technical services department on an Applied Biosystems 3130 (4-capillary) Genetic Analyzer or a 48-capillary 3730 DNA Analyzer (Both ThermoFisher Scientific, USA). Sequencing primers are listed in **Table 3**.

### Sequence analysis pipelines

#### Ion Torrent script

The fastq output file was used to align reads to the custom reference sequences, using Bowtie 2 (76). Quality control metrics were provided by BamQC (Simon Andrews, https://github.com/s-andrews/BamQC). Following this, samtools (77), bam-readcount (https://github.com/genome/bam-readcount) and the Genome Analysis Toolkit (78) packages are used to generate alignment statistics. Two different variant callers were used for comparison, VarScan 2 (79) and the Bcftools package by samtools. Bowtie 2 alignments were visualised using the Integrative Genomics Viewer (80,81).

#### MinION script

This was derived from the Ion Torrent script and is largely the same except that GraphMap (82) is used to align reads and that the following alignments are ‘cleaned up’ using Picard Tools (http://broadinstitute.github.io/picard). GraphMap contains a dedicated algorithm for aligning Oxford Nanopore data. Prior to running the MinION script, the .fast5 output was converted to .fastq using Poretools (83), and then processed using Porechop (https://github.com/rrwick/Porechop) to split the file by barcode. A BBMap script, readlength.sh (https://github.com/BioInfoTools/BBMap), was used to generate read length histograms and calculate mean/median read lengths.

For *de novo* genome assembly, Canu (84) was used to assemble MinION data. Settings were tailored to expect a small, repetitive genome. SnapGene software (from GSL Biotech; available at snapgene.com) was used to visualise the resulting genome assemblies.

### Quantitative polymerase chain reaction (qPCR)

Genomic DNA from sorted populations was subjected to qPCR. Primers used are listed in **Table 3**. Reactions were performed with PrecisionTM 2X qPCR master mix (Primerdesign) in 10 *μ*L volumes using the LightCycler® 480 System (Roche) according to the manufacturer’s instructions. The Ct values were acquired and normalised to the reference gene (*Zmynd10* exon 1) controls. The fold changes were calculated using 2^−ΔΔ*CT*^ relative quantification method.

### Animals

*Gt(ROSA)26Sor*^*tm*4(*ACTB*−*tdTomato,*−*EGFP*)^*^Luo^/J* (referred to here in the heterozygous state as FIVER) were obtained from Jackson Labs (https://www.jax.org/strain/007576)(18). *Zmynd10^em^*^1^*^Pmi^* mice were previously generated using CRISPR/Cas9 (64). Animals were maintained in an SPF environment and studies carried out in accordance with guidelines issued by the Medical Research Council in ‘Responsibility in the Use of Animals in Medical Research’ (July 1993) and licensed by the Home Office under the Animals (Scientific Procedures) Act 1986 under project license PPL P18921CDE in facilities at the University of Edinburgh (PEL 60/2605).

### Hydrodynamic tail vein injection

For NHEJ editing alone, 0.2, 2 or 20 *μ*g of SB-CRISPR-T1 plasmids were hydrodynamically co-injected (in 10% w/v physiological saline in <10s) into adult FIVER mice via the lateral tail vein with 6 *μ*g of pCMV/SB10. Mice were culled after 7 days.

For HDR editing, adult FIVER mice were given 6 *μ*g pCMV/SB10, 20 *μ*g SB-CRISPR-T1 and 20 *μ*g MC.HDR or MC.HITI. The following groups were used: N = 4 non-injected control; N = 4 SB10/CRISPR-T1 (NHEJ group); N = 3 SB10/CRISPR-T1/MC.HITI (HITI group); N = 4 SB10/CRISPR-T1/MC.HDR (HDR group). Animals were sacrificed after 7 days.

For the cancer models, adult FIVER mice were given 20 *μ*g of SB-CRISPR-T1 and 6 *μ*g pCMV/SB10, with or without 4 *μ*g of pT3-myr-AKT and 20 *μ*g pT3-NICD. N = 3 treated and N = 3 control. Animals were culled after six weeks.

### Subretinal injections

P3 FIVER animals were anaesthetised by inhalational anaesthesia (2.5% Isofluorane). Eyes were opened by cutting the fused junctional epithelium at the point where the eyelids meet. Eyes were dilated using 1% Tropicamide eye drops (Baush & Lomb). For optimal retinal view, carbomer gel was administered to the corneal surface and a 0.5 mm round coverslip placed on top. A Zeiss OPMI Lumera operating microscope was used for all procedures. Eyes were immobilised by placing traction on the rectus muscles and sclera punctured at 45° to the eye using a 34G needle (point style 12, 207434) on a 5 *μ*L Hamilton syringe (75RN, 7634-01). Needle was tunnelled subretinally towards the optic nerve prior to administration of 1.5 *μ*L of viral construct (1×108 vg, diluted in PBS) to the subretinal space. Contralateral eyes were sham injected with 1.5 *μ*L PBS to the sub retinal space as controls. Mice were sacrificed after 14 days for analysis. A 1:1:1 preparation of AAV2/5-SpCas9, AAV2/5-HDR-T1/T2 and AAV2/5-HITI was used for all experiments. Sham PBS injections were used as a control.

### Zygote injections

RNP complexes (100 ng/L Cas9 with 25 ng/*μ*L gRNA) with or without minicircle repair constructs (10 ng/*μ*L) were prepared in 0.1 TE buffer (10mM Tris-HCl, 0.1mM EDTA, pH8) and injected into fertilised outcrossed FIVER eggs via pronuclear injection and cultured for 72 hours to blastocyst stage prior to imaging.

### Cytology and histology

Animals were sacrificed 1 week post hydrodynamic tail vein injection. Livers were flushed with PBS via injection into the hepatic portal vein, then harvested and snap frozen in optimal cutting temperature compound (OCT), or fixed in 4% PFA overnight at 4°C. Following fixation, livers were incubated successively in 70% v/v, 80% v/v, 90% v/v and 100% v/v ethanol, twice in xylene and then paraffin, each for 20 min per stage with pressure, using a vacuum infiltration processor.

DAB staining was performed on 5 *μ*m paraffin liver sections. Anti-GFP (sc-8334; SantaCruz), and DSB-X biotin goat anti-chicken (D-20701; ThermoFisher Scientific, USA) antibodies were used at 1:500 and 1:2000 respectively.

OCT embedded livers were sectioned using a freezing microtome at 8 *μ*m. Sections were post fixed in 100% ethanol, washed in PBS, stained for nuclei in a 1:2500 solution of DAPI (in PBS), rinsed again in PBS and mounted using ProLong Gold antifade mounting medium (ThermoFisher Scientific, USA).

Eyes were enucleated and fixed in 4% PFA for one hour. Keratectomy and lensectomy were performed, followed by retinal dissection. Wholemount petaloid explants were prepared and explanted on slides, photoreceptor side up. Retinas were incubated in 1:1000 DRAQ5 (ThermoFisher Scientific, USA) for 5 min prior to mounting in Prolong Gold antifade mounting medium (ThermoFisher Scientific, USA).

MEFs were fixed on 6-well glass bottom dishes with 4% PFA (diluted from 16% stock in PBS; ThermoFisher Scientific, Massachusetts, US), washed with PBS, then maintained in PBS. Nuclei were stained using NucBlue Live ReadyProbes Reagent (ThermoFisher Scientific, USA). Cells were imaged using an automated pipeline (Points on a Plate PFS Surface.bin; https://tinyurl.com/yasbdqtb) using the NIS-Elements JOBS module on a Nikon widefield microscope (Nikon Instruments Europe, Netherlands).

mTECs were fixed on transwell membranes with 4% PFA (diluted from 16% stock in PBS; ThermoFisher Scientific, USA), then washed with PBS. Nuclei were stained with 1:2500 solution of DAPI (in PBS), rinsed again in PBS and mounted using ProLong Gold antifade mounting medium (ThermoFisher Scientific, USA).

### Imaging and image analysis

Fluorescent confocal images were acquired using a CFI Plan Fluor 10x 0.3NA, CFI Plan Apo VC 20x 0.75NA or CFI Plan Fluor 40x 0.75NA lens on a Nikon A1+ confocal microscope. Data were acquired using NIS-Elements AR software (Nikon Instruments Europe, Netherlands). For nuclei counting, widefield images were acquired using a CFI Plan Apo VC 20x 0.75NA lens on a Nikon Eclipse Ti microscope using NIS-Elements JOBS module in NIS-Elements AR software (Nikon Instruments Europe, Netherlands).

Retinal wholemounts were imaged using a CFI Plan Fluor 40x 0.75NA, CFI Apo Lambda S 60x 1.4NA or CFI Plan Apo 100x 1.4NA lens on an Andor Dragonfly spinning disc microscope (Oxford Instruments, UK). Data were acquired using Fusion software (Oxford Instruments, UK) and analysed using Imaris software (Oxford Instruments, UK).

DAB stained slides were imaged on a NanoZoomer XR slide scanner (Hamamatsu, Japan).

Time lapse analysis was carried out using FIJI (85) (version 2.0.0-rc-54/1.51h). Cells were tracked using the manual tracking plugin (Fabrice Cordelières, Institut Curie, Orsay, France), then mean fluorescent intensity was calculated for each time point using an automated macro.

Automated nuclei counting was carried out using a pipeline developed in QuPath (version 0.2.0-m4) (86). Total nuclei number was determined based on Hoechst staining (NucBlue Live Ready Probes Reagent; ThermoFisher Scientific, USA) using the watershed cell detection function in QuPath. The cell expansion parameter of this function was set to 1 *μ*m to create a “ring” around the nucleus, in order to sample the cytoplasm. A script was used to create a new measurement of the ratio of mean intensity of EGFP signal in the “ring” compared to that of the nucleus. This ratio measurement was used to classify all cells as having undergone HDR or not due to cells with a higher ratio having much higher mean EGFP intensity in the nucleus than cytoplasm. Cells with a ratio closer to one had either high or low mean intensity EGFP in both the nucleus and cytoplasm, more indicative of NHEJ or no editing (Classify_Ratio_Nucleus_Band_MEFs.groovy and Classify_Ratio_Nucleus_Band_mTEC.groovy; https://tinyurl.com/ycbcoopk).

### Statistics

All statistical analysis was carried out using GraphPad Prism 8 (version 8.4.1; GraphPad software, USA) as described in the text.

## Acknowledgments

We thank the IGMM technical services, FACS and Advanced Imaging facilities for support and advice. We also thank the Viral Vectors Core at the Shared University Research Facilities and the Edinburgh Wellcome Trust Clinical Research Facility Genetics Core for their technical support. We are grateful to the CBS animal facility for technical assistance and support throughout the project. We thank Chris Boyd, Ian Jackson, Wendy Bickmore, Andrew Wood and Ian Adams for helpful discussions and advice. In addition, we acknowledge Melissa Jungnickel, Lewis MacDonald, Veronica Duffy and Jennifer Brisbane for their help developing this resource. This work was supported by core funding from the MRC (MC UU 00007/14), as well as funding from MRC Confidence in Concept, Wellcome Institutional Strategic Support Fund 2 and Wellcome Seed Award (215343/Z/19/Z). Some figures were created with BioRender.com.

---

**Supplementary Table 1.**
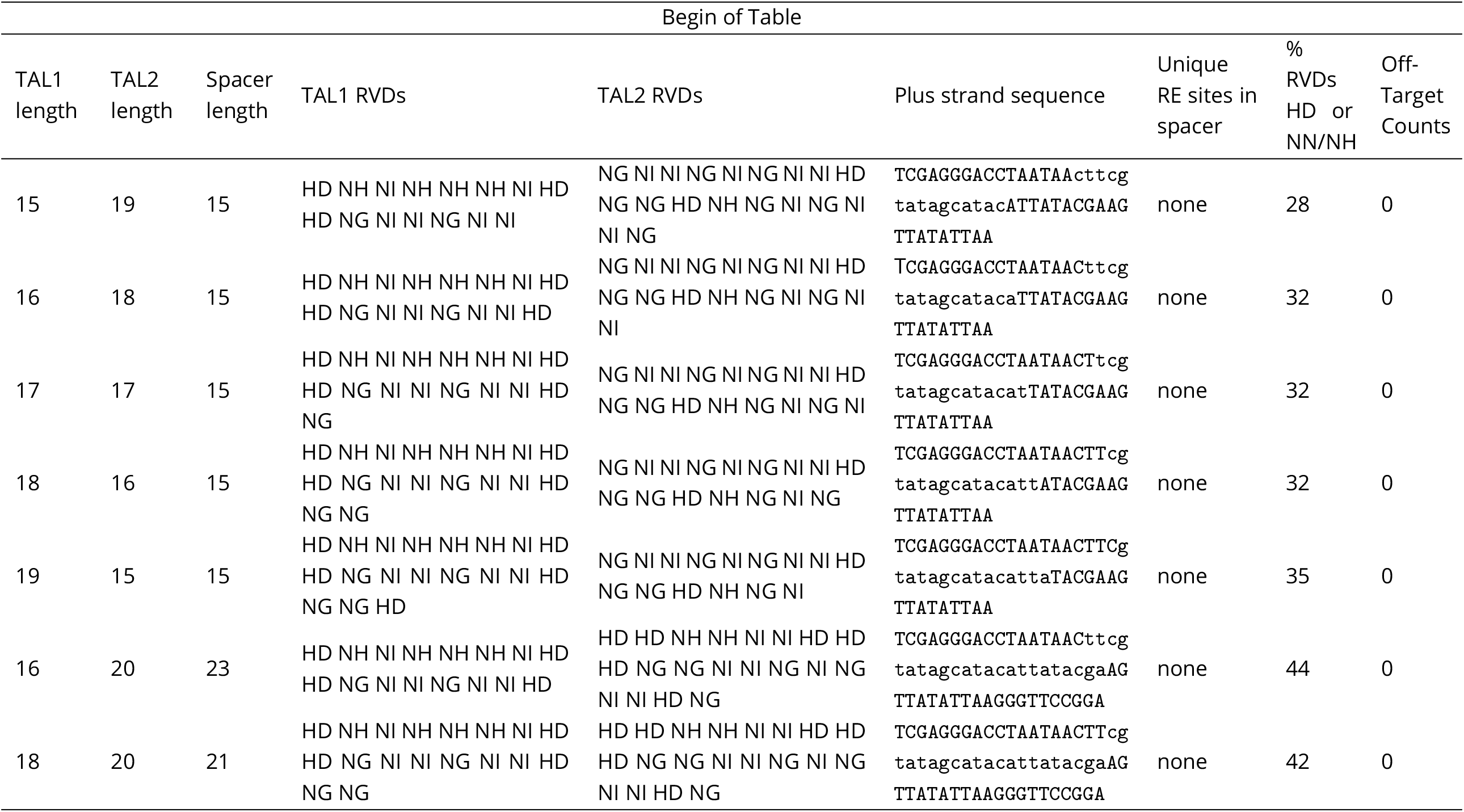
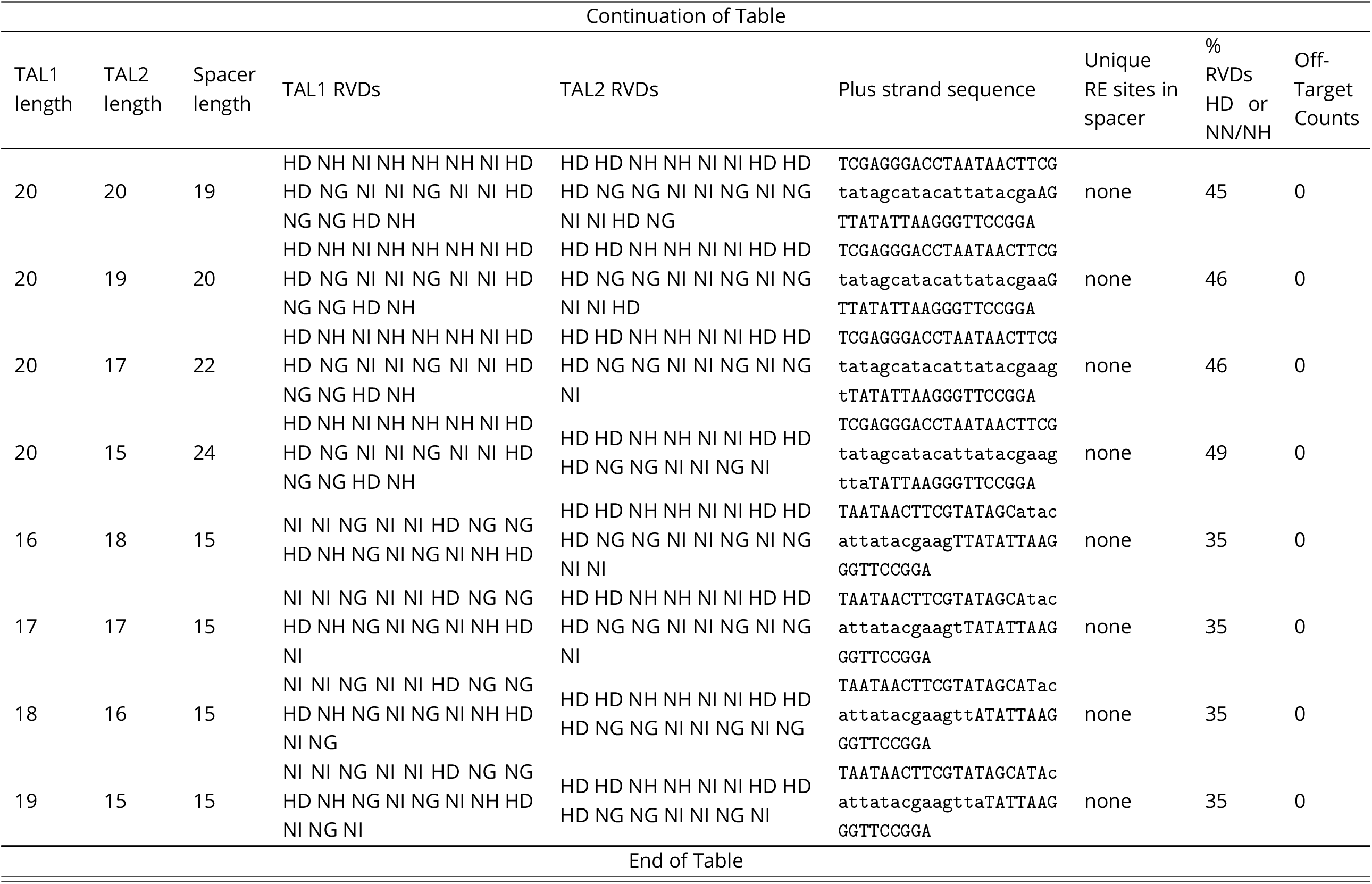
TALEN target sites within conserved region lanking tdTomato cassette. Options used: array minimum = 15; array maximum = 20; spacer minimum = 15; spacer maximum = 24 and upstream base = T. RVD = repeat variable diresidue.

## Supplementary Sequences

### MC.HDR (FIVER)

GAACAAAGGCTGCGTGCGGGGTGTGTGCGTGGGGGGGTGAGCAGGGGGTGTGGGCGCGTCGGTCGGGCTGCAACCCCCCTGCACCC CCCTCCCCGAGTTGCTGAGCACGGCCCGGCTTCGGGTGCGGGGCTCCGTACGGGGCGTGGCGCGGGGCTCGCCGTGCCGGGCGGGG GGTGGCGGCAGGTGGGGGTGCCGGGCGGGGCGGGGCCGCCTCGGGCCGGGGAGGGCTCGGGGGAGGGGCGCGGCGGCCCCCGGAGC GCCGGCGGCTGTCGAGGCGCGGCGAGCCGCAGCCATTGCCTTTTATGGTAATCGTGCGAGAGGGCGCAGGGACTTCCTTTGTCCCA AATCTGTGCGGAGCCGAAATCTGGGAGGCGCCGCCGCACCCCCTCTAGCGGGCGCGGGGCGAAGCGGTGCGGCGCCGGCAGGAAGG AAATGGGCGGGGAGGGCCTTCGTGCGTCGCCGCGCCGCCGTCCCCTTCTCCCTCTCCAGCCTCGGGGCTGTCCGCGGGGGGACGGC TGCCTTCGGGGGGGACGGGGCAGGGCGGGGTTCGGCTTCTGGCGTGTGACCGGCGGCTCTAGAGCCTCTGCTAACCATGTTCATGC CTTCTTCTTTTTCCTACAGCTCCTGGGCAACGTGCTGGTTATTGTGCTGTCTCATCATTTTGGCAAAGAATTGATTTGATACCGCG GGCCCTCGACACTAGTGAACCTCTTCGAGGGATCTAATAACTTCGTATAGCATACATTATACGAAGTTATATTAAGGGTTCCGTAC CGCCATGCCAGAGCCAGCGAAGTCTGCTCCCGCCCCGAAAAAGGGCTCCAAGAAGGCGGTGACTAAGGCGCAGAAGAAAGGCGGCA AGAAGCGCAAGCGCAGCCGCAAGGAGAGCTATTCCATCTATGTGTACAAGGTTCTGAAGCAGGTCCACCCTGACACCGGCATTTCG TCCAAGGCCATGGGCATCATGAATTCGTTTGTGAACGACATTTTCGAGCGCATCGCAGGTGAGGCTTCCCGCCTGGCGCATTACAA CAAGCGCTCGACCATCACCTCCAGGGAGATCCAGACGGCCGTGCGCCTGCTGCTGCCTGGGGAGTTGGCCAAGCACGCCGTGTCCG AGGGTACTAAGGCCATCACCAAGTACACCAGCGCTAAGGATCCACCGGTCGCCACCGTGAGCAAGGGCGAGGAGCTGTTCACCGGG GTGGTGCCCATCCTGGTCGAGCTGGACGGCGACGTAAACGGCCACAAGTTCAGCGTGTCCGGCGAGGGCGAGGGCGATGCCACCTA CGGCAAGCTGACCCTGAAGTTCATCTGCACCACCGGCAAGCTGCCCGTGCCCTGGCCCACCCTCGTGACCACCCTGACCTACGGCG TGCAGTGCTTCAGCCGCTACCCCGACCACATGAAGCAGCACGACTTCTTCAAGTCCGCCATGCCCGAAGGCTACGTCCAGGAGCGC ACCATCTTCTTCAAGGACGACGGCAACTACAAGACCCGCGCCGAGGTGAAGTTCGAGGGCGACACCCTGGTGAACCGCATCGAGCT GAAGGGCATCGACTTCAAGGAGGACGGCAACATCCTGGGGCACAAGCTGGAGTACAACTACAACAGCCACAACGTCTATATCATGG CCGACAAGCAGAAGAACGGCATCAAGGTGAACTTCAAGATCCGCCACAACATCGAGGACGGCAGCGTGCAGCTCGCCGACCACTAC CAGCAGAACACCCCCATCGGCGACGGCCCCGTGCTGCTGCCCGACAACCACTACCTGAGCACCCAGTCCGCCCTGAGCAAAGACCC CAACGAGAAGCGCGATCACATGGTCCTGCTGGAGTTCGTGACCGCCGCCGGGATCACTCTCGGCATGGACGAG

### MC.HITI (FIVER)

AGATCTGTATGCTATACGAAGTTATTAGGATCATCACCGCGGATGGGTTGCTGTGCTAGCTTGGGTGCGTTGGTTGTGGATAAGTA GCTAGACTCCAGCAACCAGTAACCTCTGCCCTTTCTCCTCCATGACAACCAGGTCCCAGGTCCCGAAAACCAAAGAAGAAGAACAT GCCAGAGCCAGCGAAGTCTGCTCCCGCCCCGAAAAAGGGCTCCAAGAAGGCGGTGACTAAGGCGCAGAAGAAAGGCGGCAAGAAGC GCAAGCGCAGCCGCAAGGAGAGCTATTCCATCTATGTGTACAAGGTTCTGAAGCAGGTCCACCCTGACACCGGCATTTCGTCCAAG GCCATGGGCATCATGAATTCGTTTGTGAACGACATTTTCGAGCGCATCGCAGGTGAGGCTTCCCGCCTGGCGCATTACAACAAGCG CTCGACCATCACCTCCAGGGAGATCCAGACGGCCGTGCGCCTGCTGCTGCCTGGGGAGTTGGCCAAGCACGCCGTGTCCGAGGGTA CTAAGGCCATCACCAAGTACACCAGCGCTAAGGATCCACCGGTCGCCACCATGAGCGAGCTGATTAAGGAGAACATGCACATGAAG CTGTACATGGAGGGCACCGTGGACAACCATCACTTCAAGTGCACATCCGAGGGCGAAGGCAAGCCCTACGAGGGCACCCAGACCAT GAGAATCAAGGTGGTCGAGGGCGGCCCTCTCCCCTTCGCCTTCGACATCCTGGCTACTAGCTTCCTCTACGGCAGCAAGACCTTCA TCAACCACACCCAGGGCATCCCCGACTTCTTCAAGCAGTCCTTCCCTGAGGGCTTCACATGGGAGAGAGTCACCACATACGAAGAC GGGGGCGTGCTGACCGCTACCCAGGACACCAGCCTCCAGGACGGCTGCCTCATCTACAACGTCAAGATCAGAGGGGTGAACTTCAC ATCCAACGGCCCTGTGATGCAGAAGAAAACACTCGGCTGGGAGGCCTTCACCGAGACGCTGTACCCCGCTGACGGCGGCCTGGAAG GCAGAAACGACATGGCCCTGAAGCTCGTGGGCGGGAGCCATCTGATCGCAAACATCAAGACCACATATAGATCCAAGAAACCCGCT AAGAACCTCAAGATGCCTGGCGTCTACTATGTGGACTACAGACTGGAAAGAATCAAGGAGGCCAACAACGAGACCTACGTCGAGCA GCACGAGGTGGCAGTGGCCAGATACTGCGACCTCCCTAGCAAACTGGGGCACAAGCTTAATTAAAGCGGCCGCTCGAGCCTCGACT GTGCCTTCTAGTTGCCAGCCATCTGTTGTTTGCCCCTCCCCCGTGCCTTCCTTGACCCTGGAAGGTGCCACTCCCACTGTCCTTTC CTAATAAAATGAGGAAATTGCATCGCATTGTCTGAGTAGGTGTCATTCTATTCTGGGGGGTGGGGTGGGGCAGGACAGCAAGGGGG AGGATTGGGAAGACAATAGCAGGCATGCTGGGGATGCGGTGGGCTCTATGGCTTCTGAGCATAGGGATCC

### MC.HITI (*Zmynd10*)

AGATCTAGCATTCACCCTGCCTGTGGAGGATCATCACCGCGGATGGGTTGCTGTGCTAGCTTGGGTGCGTTGGTTGTGGATAAGTA GCTAGACTCCAGCAACCAGTAACCTCTGCCCTTTCTCCTCCATGACAACCAGGTCCCAGGTCCCGAAAACCAAAGAAGAAGAACGA GCTGCAAAAGCAGGCGGAGATGATGGAATTTGAGATATCCCTGAAAGCCCTCTCGGTGCTTCGCTACATCACAGACTGCGTGGATA GCCTTTCCCTGAGCACACTGAACCGCATGCTCAGGACTCACAACTTGCCCTGCCTCTTGGTGGAACTGCTGGAGCACAGTCCCTGG AGCCGGCGGGTAGGAGGCAAGCTGCAGCATTTTGAGAGTGGCCGATGGCAGACGGTGGCCCCCTCAGAGCAGCAAAAGCTGAATAA ACTGGATGGGCAAGTATGGATCGCCCTGTACAATCTACTGCTCAGCCCTGAGGCCCGAGCCCGTTACTGCCTTACAAGCTTTGCCA AGGGACAGCTGCTTAAGCTTCAGGCCTTCCTCACTGACACACTACTCGACCAGTTGCCCAATCTTGCCGATCTGAAGGGTTTCCTG GCCCACCTGTCCCTGGCTGAAACCCAGCCCCCTAAGAAGGACCTAGTGTTAGAACAGATCCCAGAAATCTGGGATCGCCTGGAGAG AGAGAACAAAGGGAAATGGCAGGCTATCGCCAAGCACCAGCTTCAGCACGTATTCAGCCTCTCGGAGAAGGATCTTCGTCAACAAG CACAGAGGTGGGCTGAAACCTACAGGCTGGATGTCCTAGAGGCAGTAGCTCCGGAGAGGCCCCGCTGCGGCTACTGCAACGCAGAG GCCTCCAAGCGCTGCTCCAGATGCCAGAATGTGTGGTATTGCTGCAGGGAGTGTCAAGTCAAGCACTGGGAGAAGCACGGAAAGAC ATGTGTTCTAGCAGCCCAAGGTGACAGAGCCAAGTGAAGCGGCCGCTCGAGCCTCGAAACTTGTTTATTGCAGCTTATAATGGTTA CAAATAAAGCAATAGCATCACAAATTTCACAAATAAAGCATTTTTTTCACTGCATTCTAGTTGTGGTTTGTCCAAACTCATCAATG TATCTTATCATGTCTGGATCCTTCTGAGCATAGGGATCCCCGAATTCCGTCGACCCATGGGGGCCCGCCCCAACTGGGGTAACC

---

**Figure 1–Figure supplement 1.**
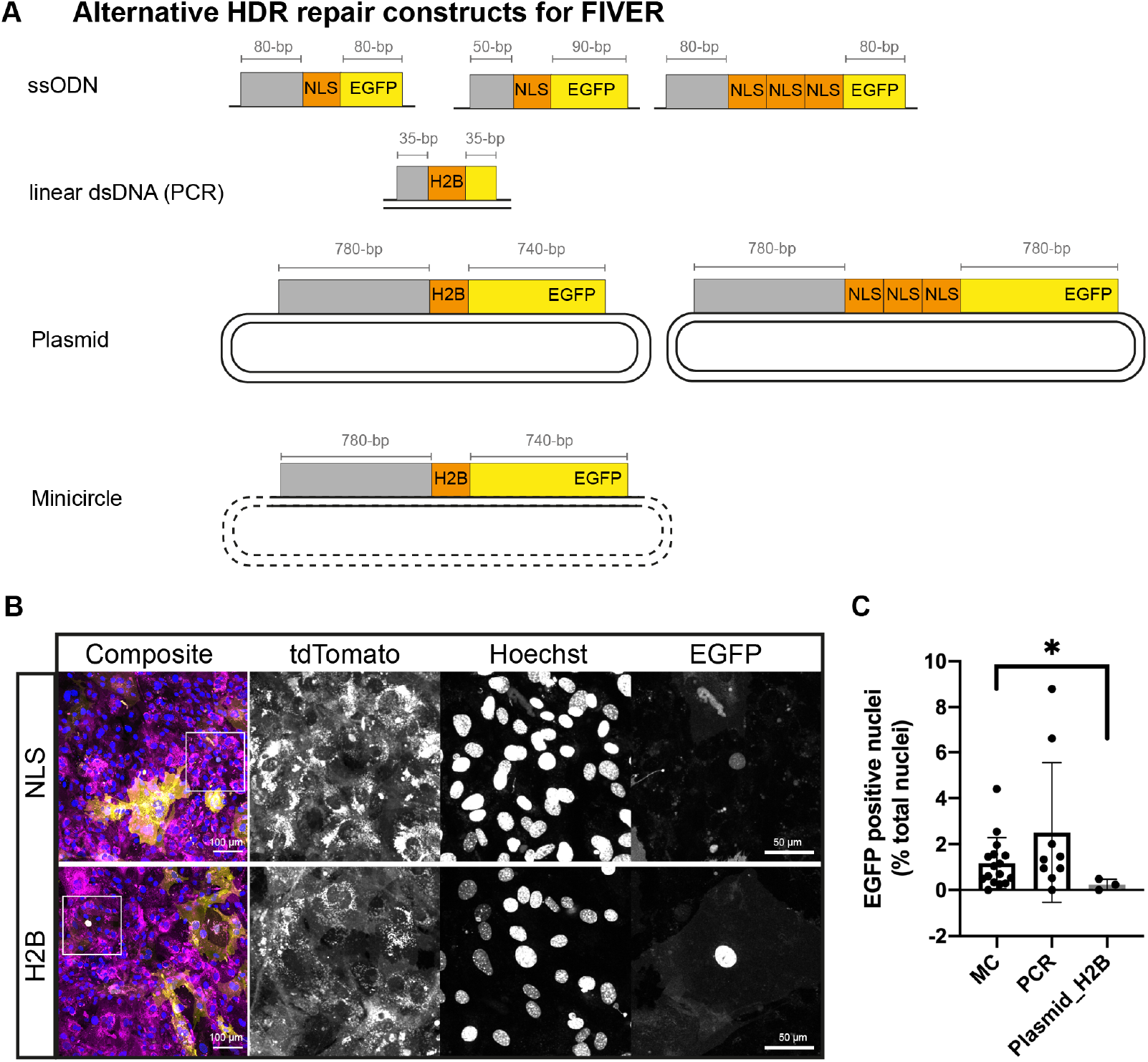
Overview of luorescent *in vivo* editing reporter (FIVER) system. (A) Schematic of alternative HDR constructs. Length of homology arms in each case indicated. Grey boxes indicate homology to sequence upstream of tdTomato, extending into the chimeric intron region. (B) Comparison of different nuclear localisation signals. Representative confocal images showing strength of nuclear signal driven by plasmid-derived 3xNLS or H2B tags. Images are maximum intensity projections of z-stacks. NLS = nuclear localisation signal, H2B = human histone H2B. (C) Assessment of HDR in FIVER MEFs after transfection with different repair constructs. HDR was determined by counting number of EGFP positive nuclei and total nuclei using an automated pipeline, n >10 cells, N ≥ 3 technical replicates. * *p* = 0.0229, one-way ANOVA with Brown-Forsythe and Welch’s multiple comparisons.

**Figure 1–Figure supplement 2.**
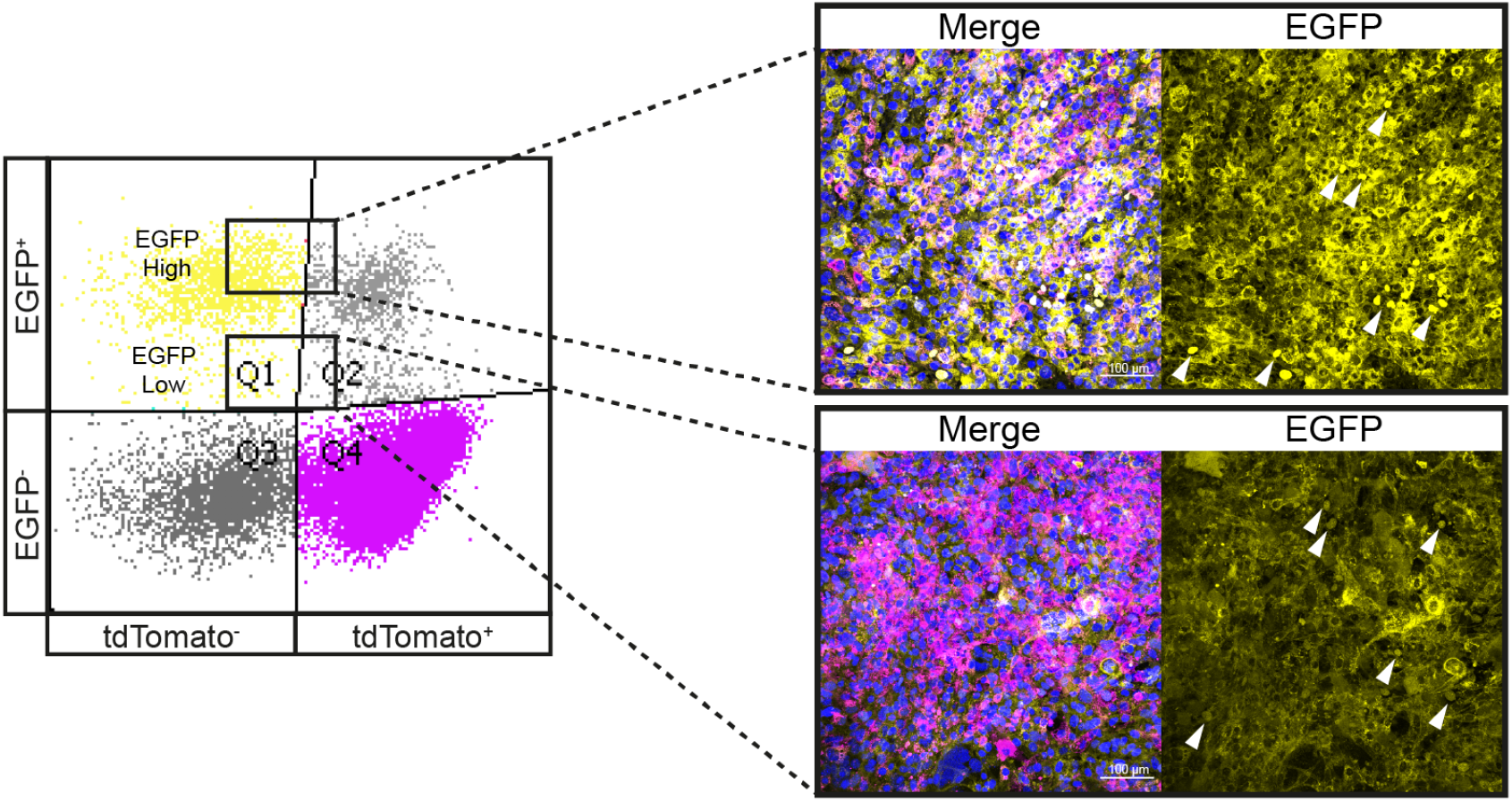
Overview of luorescent *in vivo* editing reporter (FIVER) system. Representative confocal maximum intensity projection images of sorted MEF populations. MEFs were transfected with RNPs and MC.HDR repair template. 5 days post transfection, FACS was carried out to investigate ‘high’ and ‘low’ EGFP populations for presence of nEGFP. Arrowheads indicate presence of nEGFP. Scale bar 100 *μ*m.

**Figure 2–Figure supplement 1.**
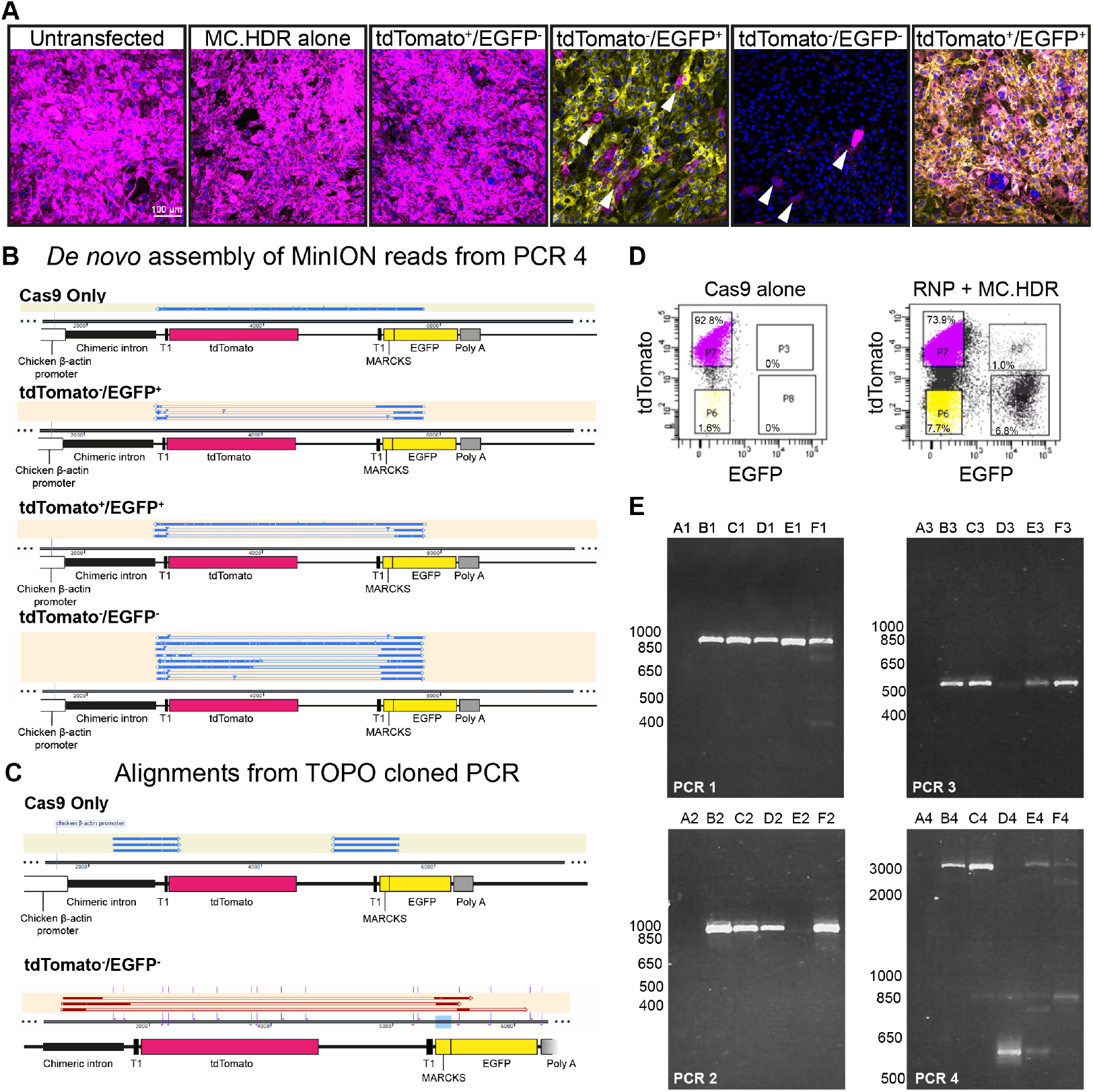
Deep sequencing confirms editing outcomes observed by FIVER. (A) Representative confocal maximum intensity projection images of edited MEF populations after FACS. Arrowheads show infiltration of tdTomato^+^ cells into other sorted populations. Scale bar 100 *μ*m. (B) Alignments for *de novo* genome assembly of MinION reads from PCR 4. Assembled sequences are ordered based on the number of reads from which they were generated; assembled sequences generated from the greatest number of reads are uppermost. (C) Reads from TOPO cloning following amplification with P7-P8 (PCR 5) and P7-P9 (PCR 6) were aligned to reference sequences. Example alignments for PCR 6 are presented. (D) FACS plots illustrating gating used to sort each population for sequencing: tdTomato^+^/EGFP^−^ (400,000), tdTomato^−^/EGFP^+^ (20,000), tdTomato^−^/EGFP^−^ (20,000) and tdTomato^+^/EGFP^+^ (3,000). (E) Purified PCR products were analysed by agarose gel electrophoresis prior to sequencing. A = no template control. B = Cas9 only, tdTomato^+^, C = tdTomato^+^/EGFP^−^, D = tdTomato^−^/EGFP^+^, E = tdTomato^+^/EGFP^+^ and F = tdTomato^−^/EGFP^−^. Sizes are indicated in bp.

**Figure 4–Figure supplement 1.**
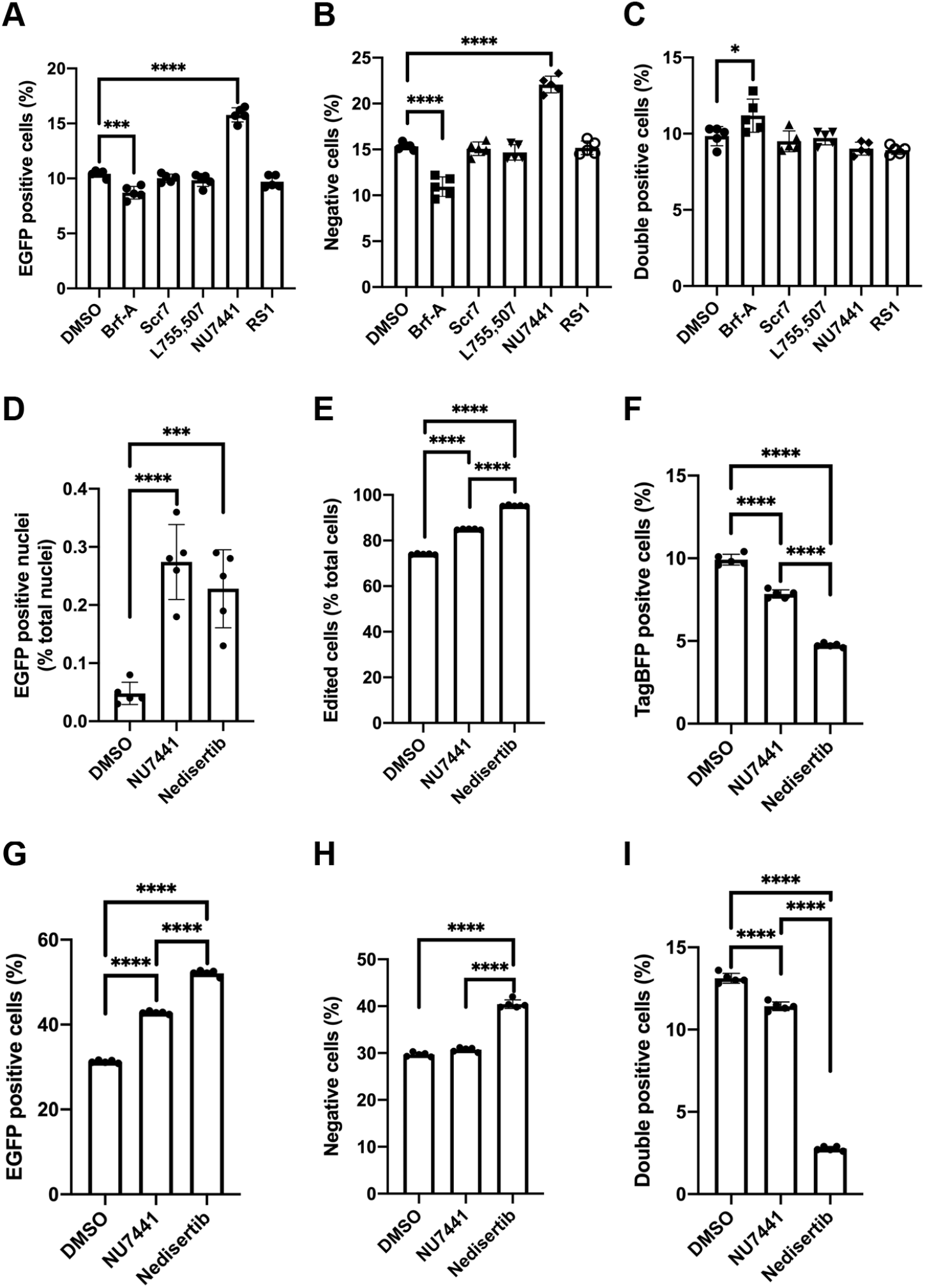
Small molecule modulators of genome editing outcome. Editing outcomes were determined by flow cytometry after treatment with Brf-A (0.1 *μ*M), Scr7 (0.1 *μ*M), L755,507 (5 *μ*M), NU7441 (2 *μ*M) or RS1 (10 *μ*M) for 24 hours. (A) Total tdTomato^−^/EGFP^+^ cells, n = 60,000 cells, N = 5 technical replicates. (B) Total tdTomato^−^/EGFP^−^ cells, n = 60,000 cells, N = 5. (C) Total tdTomato^+^/EGPF^+^ cells, n = 60,000 cells, N = 5. Next, cells were treated with NU7441 (2 *μ*M) or Nedisertib (2 *μ*M) for 24 hours and editing outcomes determined by flow cytometry. (D) EGFP positive nuclei, determined by widefield microscopy, n > 10,000 cells, N = 5. (E) Total edited cells, determined by flow cytometry, n = 100,000 cells, N = 5. (F) Total TagBFP^+^ cells, determined by flow cytometry, n = 100,000 cells, N = 5. (G) Total tdTomato^−^/EGFP^+^ cells, n = 100,000 cells, N = 5. (H) Total tdTomato^−^/EGFP^−^ cells, n = 100,000 cells, N = 5. (I) Total tdTomato^+^/EGPF^+^ cells, n = 100,000 cells, N = 5. Significance was tested using one-way ANOVA and Dunnett’s (for comparison to DMSO control) or Tukey’s (for all comparisons) multiple comparisons, 0.0021 < *p* < 0.05 = *, 0.0002 < *p* < 0.0021 = **, 0.0001 < *p* < 0.0002 = ***, *p* < 0.0001 = ****.

**Figure 4–Figure supplement 2.**
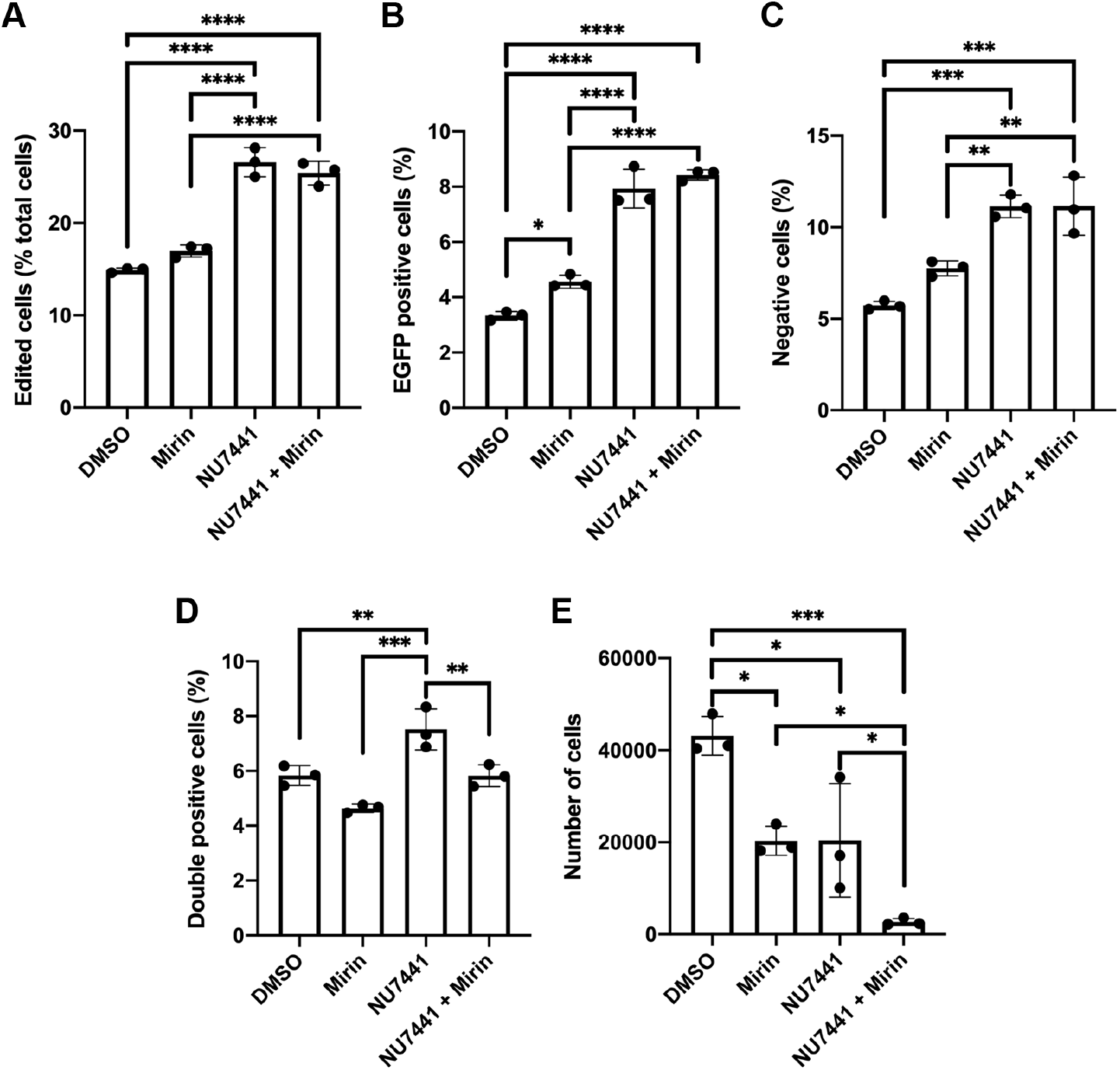
Small molecule modulators of genome editing outcome. Editing outcomes were determined by flow cytometry 72 hours post transfection, following 24-hour treatment with mirin (50 *μ*M) and NU7441 (2 *μ*M), alone or in combination, immediately after transfection. (A) Total edited cells, n > 2,000 cells, N = 3 technical replicates. (B) Total tdTomato^−^/EGFP^+^ cells, n > 2,000 cells, N = 3 technical replicates. (C) Total tdTomato^−^/EGFP^−^ cells, n > 2,000 cells, N = 3 technical replicates. (D) Total tdTomato^−^/EGFP^+^ cells, n > 2,000 cells, N = 3 technical replicates. (E) Total cells sorted in 2 minutes, n > 2,000 cells, N = 3 technical replicates. Significance was tested using one-way ANOVA and or Tukey’s multiple comparisons, 0.0021 < *p* < 0.05 = *, 0.0002 < *p* < 0.0021 = **, 0.0001 < *p* < 0.0002 = ***, *p* < 0.0001 = ****.

**Figure 5–Figure supplement 1.**
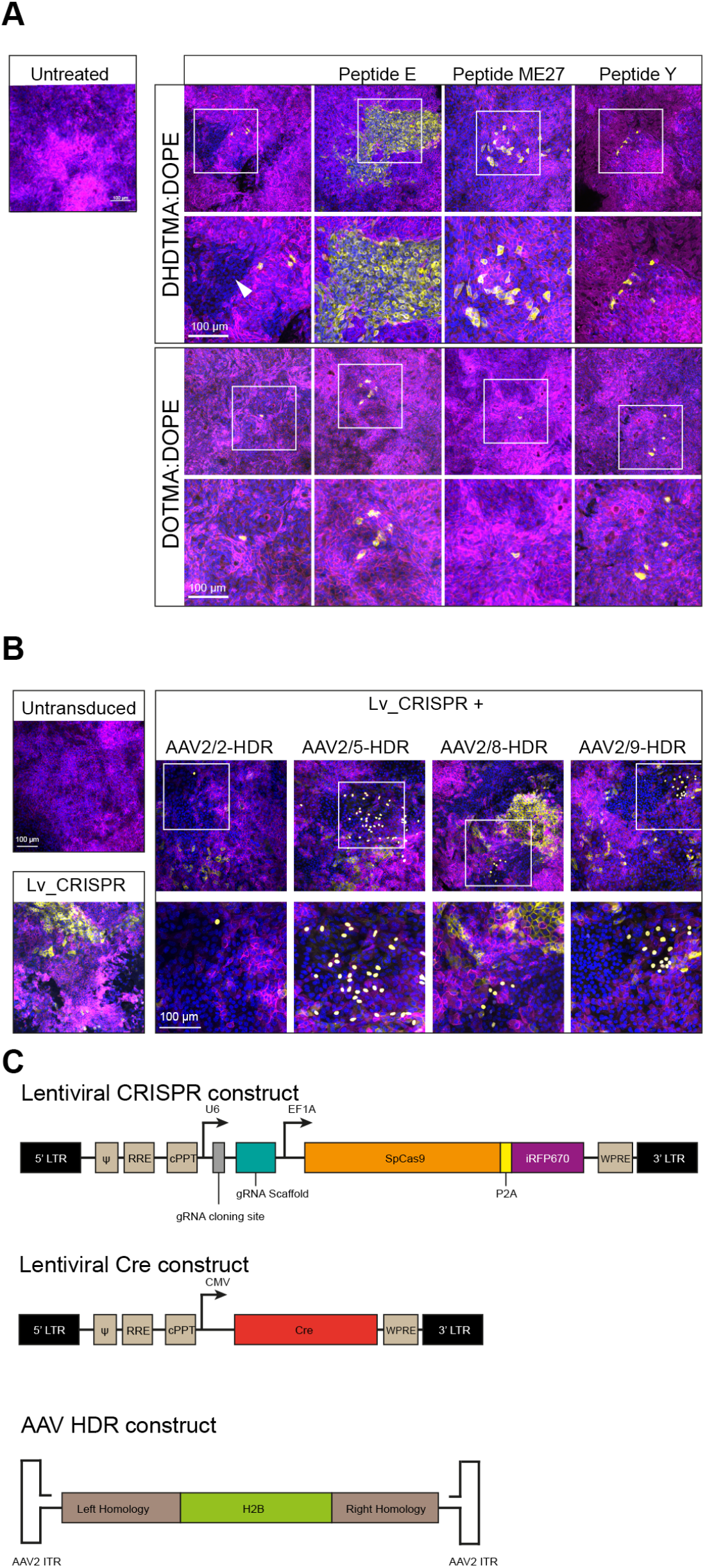
FIVER allows establishment of disease-relevant primary cultures and organoids. (A) Representative confocal images of mTECs treated with lipid nanoparticles containing Cas9-T1 RNPs and MC.HDR. NHEJ editing indicated by mEGFP fluorescence or loss of mtdTomato (arrowhead). Nuclei visualised with DAPI. (B) Representative confocal images following transduction of mTECs with different AAV serotypes in conjunction with lentiviral delivered CRISPR machinery. Nuclei visualised with DAPI. (C) Viral constructs for delivery of CRISPR machinery and HDR construct. Lv-Cre was used as a positive control for ductal liver organoid delivery, see **Figure 5C**. All images are maximum intensity projections of z-stacks.

## References

1. Schacker M, Seimetz D. From fiction to science: clinical potentials and regulatory considerations of gene editing. Clin Transl Med. 2019 Oct 21;8(1):27.

2. Mullard A. Gene-editing pipeline takes off. Nat Rev Drug Discov. 2020 May 15;

3. Giannoukos G, Ciulla DM, Marco E, Abdulkerim HS, Barrera LA, Bothmer A, et al. UDiTaSTM, a genome editing detection method for indels and genome rearrangements. BMC Genomics. 2018 Mar 21;19(1):212.

4. Nelson CE, Wu Y, Gemberling MP, Oliver ML, Waller MA, Bohning JD, et al. Long-term evaluation of AAV-CRISPR genome editing for Duchenne muscular dystrophy. Nat Med. 2019 Feb 18;25(3):427–32.

5. Kim D, Kim J, Hur JK, Been KW, Yoon S-H, Kim J-S. Genome-wide analysis reveals specificities of Cpf1 endonucleases in human cells. Nat Biotechnol. 2016 Jun 6;34(8):863–8.

6. Akcakaya P, Bobbin ML, Guo JA, Malagon-Lopez J, Clement K, Garcia SP, et al. In vivo CRISPR editing with no detectable genome-wide off-target mutations. Nature. 2018 Sep 12;561(7723): 416–9.

7. Pinello L, Canver MC, Hoban MD, Orkin SH, Kohn DB, Bauer DE, et al. Analyzing CRISPR genome-editing experiments with CRISPResso. Nat Biotechnol. 2016 Jul 12;34(7):695–7.

8. Hendel A, Kildebeck EJ, Fine EJ, Clark J, Punjya N, Sebastiano V, et al. Quantifying genome-editing outcomes at endogenous loci with SMRT sequencing. Cell Rep. 2014 Apr 10;7(1): 293–305.

9. Ceccaldi R, Rondinelli B, D’Andrea AD. Repair Pathway Choices and Consequences at the Double-Strand Break. Trends Cell Biol. 2016 Jan;26(1):52–64.

10. Certo MT, Ryu BY, Annis JE, Garibov M, Jarjour J, Rawlings DJ, et al. Tracking genome engineering outcome at individual DNA breakpoints. Nat Methods. 2011 Jul 10;8(8):671–6.

11. Kuhar R, Gwiazda KS, Humbert O, Mandt T, Pangallo J, Brault M, et al. Novel fluorescent genome editing reporters for monitoring DNA repair pathway utilization at endonuclease-induced breaks. Nucleic Acids Res. 2014 Jan;42(1):e4.

12. Yang F, Liu C, Chen D, Tu M, Xie H, Sun H, et al. CRISPR/Cas9-loxP-Mediated Gene Editing as a Novel Site-Specific Genetic Manipulation Tool. Mol Ther Nucleic Acids. 2017 Jun 16;7:378–86.

13. Xie H, Tang L, He X, Liu X, Zhou C, Liu J, et al. SaCas9 Requires 5’-NNGRRT-3’ PAM for Sufficient Cleavage and Possesses Higher Cleavage Activity than SpCas9 or FnCpf1 in Human Cells. Biotechnol J. 2018 Apr;13(4):e1700561.

14. Zhang H, Zhou Y, Wang Y, Zhao Y, Qiu Y, Zhang X, et al. A surrogate reporter system for multiplexable evaluation of CRISPR/Cas9 in targeted mutagenesis. Sci Rep. 2018 Jan 18;8(1):1042.

15. Alapati D, Zacharias WJ, Hartman HA, Rossidis AC, Stratigis JD, Ahn NJ, et al. In utero gene editing for monogenic lung disease. Sci Transl Med. 2019 Apr 17;11(488).

16. Glaser A, McColl B, Vadolas J. GFP to BFP Conversion: A Versatile Assay for the Quantification of CRISPR/Cas9-mediated Genome Editing. Mol Ther Nucleic Acids. 2016 Jul 12;5(7):e334.

17. Richardson CD, Ray GJ, DeWitt MA, Curie GL, Corn JE. Enhancing homology-directed genome editing by catalytically active and inactive CRISPR-Cas9 using asymmetric donor DNA. Nat Biotechnol. 2016 Mar;34(3):339–44.

18. Suzuki K, Tsunekawa Y, Hernandez-Benitez R, Wu J, Zhu J, Kim EJ, et al. In vivo genome editing via CRISPR/Cas9 mediated homology-independent targeted integration. Nature. 2016 Dec 1;540(7631):144–9.

19. Muzumdar MD, Tasic B, Miyamichi K, Li L, Luo L. A global double-fluorescent Cre reporter mouse. Genesis. 2007 Sep;45(9):593–605.

20. Iyama T, Wilson DM. DNA repair mechanisms in dividing and non-dividing cells. DNA Repair (Amst). 2013 Aug;12(8):620–36.

21. Mao Z, Bozzella M, Seluanov A, Gorbunova V. DNA repair by nonhomologous end joining and homologous recombination during cell cycle in human cells. Cell Cycle. 2008 Sep 15;7(18): 2902–6.

22. Lino CA, Harper JC, Carney JP, Timlin JA. Delivering CRISPR: a review of the challenges and approaches. Drug Deliv. 2018 Nov;25(1):1234–57.

23. Gracey Maniar LE, Maniar JM, Chen Z-Y, Lu J, Fire AZ, Kay MA. Minicircle DNA vectors achieve sustained expression reflected by active chromatin and transcriptional level. Mol Ther. 2013 Jan;21(1):131–8.

24. Gaspar V, de Melo-Diogo D, Costa E, Moreira A, Queiroz J, Pichon C, et al. Minicircle DNA vectors for gene therapy: advances and applications. Expert Opin Biol Ther. 2015 Mar;15(3): 353–79.

25. Maruyama T, Dougan SK, Truttmann MC, Bilate AM, Ingram JR, Ploegh HL. Increasing the efficiency of precise genome editing with CRISPR-Cas9 by inhibition of nonhomologous end joining. Nat Biotechnol. 2015 May;33(5):538–42.

26. Mali P, Yang L, Esvelt KM, Aach J, Guell M, DiCarlo JE, et al. RNA-guided human genome engineering via Cas9. Science. 2013 Feb 15;339(6121):823–6.

27. Yu C, Liu Y, Ma T, Liu K, Xu S, Zhang Y, et al. Small molecules enhance CRISPR genome editing in pluripotent stem cells. Cell Stem Cell. 2015 Feb 5;16(2):142–7.

28. Orthwein A, Noordermeer SM, Wilson MD, Landry S, Enchev RI, Sherker A, et al. A mechanism for the suppression of homologous recombination in G1 cells. Nature. 2015 Dec 17;528(7582):422–6.

29. Srivastava M, Nambiar M, Sharma S, Karki SS, Goldsmith G, Hegde M, et al. An inhibitor of nonhomologous end-joining abrogates double-strand break repair and impedes cancer progression. Cell. 2012 Dec 21;151(7):1474–87.

30. Robert F, Barbeau M, Éthier S, Dostie J, Pelletier J. Pharmacological inhibition of DNA-PK stimulates Cas9-mediated genome editing. Genome Med. 2015 Aug 27;7:93.

31. Leahy JJJ, Golding BT, Griffin RJ, Hardcastle IR, Richardson C, Rigoreau L, et al. Identification of a highly potent and selective DNA-dependent protein kinase (DNA-PK) inhibitor (NU7441) by screening of chromenone libraries. Bioorg Med Chem Lett. 2004 Dec 20;14(24):6083–7.

32. Pinder J, Salsman J, Dellaire G. Nuclear domain “knock-in” screen for the evaluation and identification of small molecule enhancers of CRISPR-based genome editing. Nucleic Acids Res. 2015 Oct 30;43(19):9379–92.

33. Jayathilaka K, Sheridan SD, Bold TD, Bochenska K, Logan HL, Weichselbaum RR, et al. A chemical compound that stimulates the human homologous recombination protein RAD51. Proc Natl Acad Sci USA. 2008 Oct 14;105(41):15848–53.

34. Song J, Yang D, Xu J, Zhu T, Chen YE, Zhang J. RS-1 enhances CRISPR/Cas9- and TALEN-mediated knock-in efficiency. Nat Commun. 2016 Jan 28;7:10548.

35. Parmee ER, Ok HO, Candelore MR, Tota L, Deng L, Strader CD, et al. Discovery of L-755,507: A subnanomolar human *β*3 adrenergic receptor agonist. Bioorg Med Chem Lett. 1998 May;8(9): 1107–12.

36. Benabdi S, Peurois F, Nawrotek A, Chikireddy J, Cañeque T, Yamori T, et al. Family-wide Analysis of the Inhibition of Arf Guanine Nucleotide Exchange Factors with Small Molecules: Evidence of Unique Inhibitory Profiles. Biochemistry. 2017 Sep 26;56(38):5125–33.

37. Riesenberg S, Chintalapati M, Macak D, Kanis P, Maricic T, Pääbo S. Simultaneous precise editing of multiple genes in human cells. Nucleic Acids Res. 2019 Nov 4;47(19):e116.

38. Montoro DT, Haber AL, Biton M, Vinarsky V, Lin B, Birket SE, et al. A revised airway epithelial hierarchy includes CFTR-expressing ionocytes. Nature. 2018 Aug 1;560(7718):319–24.

39. Writer M, Hurley CA, Sarkar S, Copeman DM, Wong JB, Odlyha M, et al. Analysis and optimization of the cationic lipid component of a lipid/peptide vector formulation for enhanced transfection in vitro and in vivo. J Liposome Res. 2006;16(4):373–89.

40. Writer MJ, Marshall B, Pilkington-Miksa MA, Barker SE, Jacobsen M, Kritz A, et al. Targeted gene delivery to human airway epithelial cells with synthetic vectors incorporating novel targeting peptides selected by phage display. J Drug Target. 2004 May;12(4):185–93.

41. Russell DW, Hirata RK. Human gene targeting by viral vectors. Nat Genet. 1998 Apr;18(4): 325–30.

42. Gaj T, Staahl BT, Rodrigues GMC, Limsirichai P, Ekman FK, Doudna JA, et al. Targeted gene knock-in by homology-directed genome editing using Cas9 ribonucleoprotein and AAV donor delivery. Nucleic Acids Res. 2017 Jun 20;45(11):e98.

43. Payne JG, Takahashi A, Higgins MI, Porter EL, Suki B, Balazs A, et al. Multilineage transduction of resident lung cells in vivo by AAV2/8 for *α*1-antitrypsin gene therapy. Mol Ther Methods Clin Dev. 2016 Jun 29;3:16042.

44. Bell CL, Vandenberghe LH, Bell P, Limberis MP, Gao G-P, Van Vliet K, et al. The AAV9 receptor and its modification to improve in vivo lung gene transfer in mice. J Clin Invest. 2011 Jun;121(6):2427–35.

45. Asokan A, Schaffer DV, Samulski RJ. The AAV vector toolkit: poised at the clinical crossroads. Mol Ther. 2012 Apr;20(4):699–708.

46. Wilson JM. Adeno-associated Virus and Lentivirus Pseudotypes for Lung-directed Gene Therapy. Proc Am Thorac Soc. 2004 Dec 1;1(4):309–14.

47. Nantasanti S, de Bruin A, Rothuizen J, Penning LC, Schotanus BA. Concise review: organoids are a powerful tool for the study of liver disease and personalized treatment design in humans and animals. Stem Cells Transl Med. 2016 Mar;5(3):325–30.

48. Gu B, Posfai E, Rossant J. Efficient generation of targeted large insertions by microinjection into two-cell-stage mouse embryos. Nat Biotechnol. 2018 Jun 11;36(7):632–7.

49. Ciemerych MA, Sicinski P. Cell cycle in mouse development. Oncogene. 2005 Apr 18;24(17): 2877–98.

50. Yin H, Xue W, Chen S, Bogorad RL, Benedetti E, Grompe M, et al. Genome editing with Cas9 in adult mice corrects a disease mutation and phenotype. Nat Biotechnol. 2014 Jun;32(6):551–3.

51. Weber J, Öllinger R, Friedrich M, Ehmer U, Barenboim M, Steiger K, et al. CRISPR/Cas9 somatic multiplex-mutagenesis for high-throughput functional cancer genomics in mice. Proc Natl Acad Sci USA. 2015 Nov 10;112(45):13982–7.

52. Mitchell RS, Beitzel BF, Schroder ARW, Shinn P, Chen H, Berry CC, et al. Retroviral DNA integration: ASLV, HIV, and MLV show distinct target site preferences. PLoS Biol. 2004 Aug 17;2(8):E234.

53. Yant SR, Wu X, Huang Y, Garrison B, Burgess SM, Kay MA. High-resolution genome-wide mapping of transposon integration in mammals. Mol Cell Biol. 2005 Mar;25(6):2085–94.

54. Schröder ARW, Shinn P, Chen H, Berry C, Ecker JR, Bushman F. HIV-1 integration in the human genome favors active genes and local hotspots. Cell. 2002 Aug 23;110(4):521–9.

55. Ivics Z, Hackett PB, Plasterk RH, Izsvák Z. Molecular reconstruction of Sleeping Beauty, a Tc1-like transposon from fish, and its transposition in human cells. Cell. 1997 Nov 14;91(4):501–10.

56. Fan B, Malato Y, Calvisi DF, Naqvi S, Razumilava N, Ribback S, et al. Cholangiocarcinomas can originate from hepatocytes in mice. J Clin Invest. 2012 Aug;122(8):2911–5.

57. Maeder ML, Stefanidakis M, Wilson CJ, Baral R, Barrera LA, Bounoutas GS, et al. Development of a gene-editing approach to restore vision loss in Leber congenital amaurosis type 10. Nat Med. 2019 Jan 21;25(2):229–33.

58. Hampton T. With first CRISPR trials, gene editing moves toward the clinic. JAMA. 2020 Apr 8;

59. Kallimasioti-Pazi EM, Thelakkad Chathoth K, Taylor GC, Meynert A, Ballinger T, Kelder MJE, et al. Heterochromatin delays CRISPR-Cas9 mutagenesis but does not influence the outcome of mutagenic DNA repair. PLoS Biol. 2018 Dec 12;16(12):e2005595.

60. Chen X, Liu J, Janssen JM, Gonçalves MAFV. The Chromatin Structure Differentially Impacts High-Specificity CRISPR-Cas9 Nuclease Strategies. Mol Ther Nucleic Acids. 2017 Sep 15;8: 558–63.

61. Chen X, Rinsma M, Janssen JM, Liu J, Maggio I, Gonçalves MAFV. Probing the impact of chromatin conformation on genome editing tools. Nucleic Acids Res. 2016 Jul 27;44(13):6482–92.

62. Jensen KT, Fløe L, Petersen TS, Huang J, Xu F, Bolund L, et al. Chromatin accessibility and guide sequence secondary structure affect CRISPR-Cas9 gene editing efficiency. FEBS Lett. 2017 Jun 28;591(13):1892–901.

63. Schep R, Brinkman EK, Leemans C, Vergara X, Morris B, van Schaik T, et al. Impact of chromatin context on Cas9-induced DNA double-strand break repair pathway balance. BioRxiv. 2020 May 5;

64. Mali GR, Yeyati PL, Mizuno S, Dodd DO, Tennant PA, Keighren MA, et al. ZMYND10 functions in a chaperone relay during axonemal dynein assembly. Elife. 2018 Jun 19;7.

65. Yeh CD, Richardson CD, Corn JE. Advances in genome editing through control of DNA repair pathways. Nat Cell Biol. 2019 Dec 2;21(12):1468–78.

66. Wang J, Zhang C, Feng B. The rapidly advancing Class 2 CRISPR-Cas technologies: A customizable toolbox for molecular manipulations. J Cell Mol Med. 2020 Mar;24(6):3256–70.

67. Jinek M, Chylinski K, Fonfara I, Hauer M, Doudna JA, Charpentier E. A programmable dual-RNA-guided DNA endonuclease in adaptive bacterial immunity. Science. 2012 Aug 17;337(6096): 816–21.

68. Zetsche B, Gootenberg JS, Abudayyeh OO, Slaymaker IM, Makarova KS, Essletzbichler P, et al. Cpf1 is a single RNA-guided endonuclease of a class 2 CRISPR-Cas system. Cell. 2015 Oct 22;163(3):759–71.

69. Owens DDG, Caulder A, Frontera V, Harman JR, Allan AJ, Bucakci A, et al. Microhomologies are prevalent at Cas9-induced larger deletions. Nucleic Acids Res. 2019 Aug 22;47(14):7402–17.

70. Sharma S, Javadekar SM, Pandey M, Srivastava M, Kumari R, Raghavan SC. Homology and enzymatic requirements of microhomology-dependent alternative end joining. Cell Death Dis. 2015 Mar 19;6:e1697.

71. Ohmori T, Nagao Y, Mizukami H, Sakata A, Muramatsu S-I, Ozawa K, et al. CRISPR/Cas9-mediated genome editing via postnatal administration of AAV vector cures haemophilia B mice. Sci Rep. 2017 Jun 23;7(1):4159.

72. Chou S, Yang P, Ban Q, Yang Y, Wang M, Chien C, et al. Dual supramolecular nanoparticle vectors enable crispr/cas9-mediated knockin of retinoschisin 1 gene—a potential nonviral therapeutic solution for x-linked juvenile retinoschisis. Adv Sci. 2020 Apr 16;1903432.

73. Wang L, Li M-Y, Qu C, Miao W-Y, Yin Q, Liao J, et al. CRISPR-Cas9-mediated genome editing in one blastomere of two-cell embryos reveals a novel Tet3 function in regulating neocortical development. Cell Res. 2017 Jun;27(6):815–29.

74. Lang JF, Toulmin SA, Brida KL, Eisenlohr LC, Davidson BL. Standard screening methods under-report AAV-mediated transduction and gene editing. Nat Commun. 2019 Jul 30;10(1):3415.

75. Vladar EK, Brody SL. Analysis of ciliogenesis in primary culture mouse tracheal epithelial cells. Meth Enzymol. 2013;525:285–309.

76. Langmead B, Salzberg SL. Fast gapped-read alignment with Bowtie 2. Nat Methods. 2012 Mar 4;9(4):357–9.

77. Li H, Handsaker B, Wysoker A, Fennell T, Ruan J, Homer N, et al. The Sequence Alignment/Map format and SAMtools. Bioinformatics. 2009 Aug 15;25(16):2078–9.

78. McKenna A, Hanna M, Banks E, Sivachenko A, Cibulskis K, Kernytsky A, et al. The Genome Analysis Toolkit: a MapReduce framework for analyzing next-generation DNA sequencing data. Genome Res. 2010 Sep;20(9):1297–303.

79. Koboldt DC, Zhang Q, Larson DE, Shen D, McLellan MD, Lin L, et al. VarScan 2: somatic mutation and copy number alteration discovery in cancer by exome sequencing. Genome Res. 2012 Mar;22(3):568–76.

80. Robinson JT, Thorvaldsdóttir H, Winckler W, Guttman M, Lander ES, Getz G, et al. Integrative genomics viewer. Nat Biotechnol. 2011 Jan;29(1):24–6.

81. Thorvaldsdóttir H, Robinson JT, Mesirov JP. Integrative Genomics Viewer (IGV): high-performance genomics data visualization and exploration. Brief Bioinformatics. 2013 Mar;14(2):178–92.

82. Sović I, Šikić M, Wilm A, Fenlon SN, Chen S, Nagarajan N. Fast and sensitive mapping of nanopore sequencing reads with GraphMap. Nat Commun. 2016 Apr 15;7:11307.

83. Loman NJ, Quinlan AR. Poretools: a toolkit for analyzing nanopore sequence data. Bioinformatics. 2014 Dec 1;30(23):3399–401.

84. Koren S, Walenz BP, Berlin K, Miller JR, Bergman NH, Phillippy AM. Canu: scalable and accurate long-read assembly via adaptive k-mer weighting and repeat separation. Genome Res. 2017 Mar 15;27(5):722–36.

85. Schindelin J, Arganda-Carreras I, Frise E, Kaynig V, Longair M, Pietzsch T, et al. Fiji: an open-source platform for biological-image analysis. Nat Methods. 2012 Jun 28;9(7):676–82.

86. Bankhead P, Loughrey MB, Fernández JA, Dombrowski Y, McArt DG, Dunne PD, et al. QuPath: Open source software for digital pathology image analysis. Sci Rep. 2017 Dec 4;7(1):16878.

